# Ligand-induced CaMKIIα hub Trp403 flip, hub domain stacking and kinase inhibition

**DOI:** 10.1101/2024.03.26.586665

**Authors:** Dilip Narayanan, Anne Sofie G. Larsen, Stine Juul Gauger, Ruth Adafia, Rikke Bartschick Hammershøi, Louise Hamborg, Jesper Bruus-Jensen, Nane Griem-Krey, Christine L. Gee, Bente Frølund, Margaret M. Stratton, John Kuriyan, Jette Sandholm Kastrup, Annette E. Langkilde, Petrine Wellendorph, Sara M. Ø. Solbak

## Abstract

γ-Hydroxybutyric acid (GHB) analogs are small molecules that bind competitively to a specific cavity in the oligomeric CaMKIIα hub domain. Binding affects conformation and stability of the hub domain, which may explain the neuroprotective action of some of these compounds. Here, we describe molecular details of interaction of the larger-type GHB analog 2-(6-(4-chlorophenyl)imidazo[1,2-b]pyridazine-2-yl)acetic acid (PIPA). Like smaller-type analogs, PIPA binding to the CaMKIIα hub domain promoted thermal stability. PIPA additionally inhibited CaMKIIα kinase activity by reducing CaM sensitivity. A high-resolution X-ray crystal structure of a stabilized CaMKIIα (6x mutant) hub construct revealed details of the binding mode of PIPA, which involved outward placement of tryptophan 403 (Trp403), a central residue in a flexible loop close to the upper hub cavity. Small-angle X-ray scattering (SAXS) solution structures and mass photometry of the CaMKIIα wildtype hub domain in the presence of PIPA revealed a high degree of ordered self-association (stacks of CaMKIIα hub domains). This stacking neither occurred with the smaller compound 3-hydroxycyclopent-1-enecarboxylic acid (HOCPCA), nor when Trp403 was replaced with leucine (W403L). Additionally, CaMKIIα W403L hub was stabilized to a larger extent by PIPA compared to CaMKIIα hub wildtype, indicating that loop flexibility is important for holoenzyme stability. Thus, we propose that ligand-induced outward placement of Trp403 by PIPA, which promotes an unforeseen mechanism of hub domain stacking, may be involved in the observed reduction in CaMKIIα kinase activity. Altogether, this sheds new light on allosteric regulation of CaMKIIα activity via the hub domain.

## Introduction

Recently, the calcium/calmodulin-dependent protein kinase II subtype alpha (CaMKIIα) was discovered as the long-sought-after high-affinity neuronal target for γ-hydroxybutyric acid (GHB; Fig. 1A) ^1, 2^. This was surprising since GHB is a natural metabolite of γ-aminobutyric acid (GABA) involved in inhibitory neurotransmission. Intriguingly, GHB and synthetic small-molecule analogs were found to bind to a novel pocket in the hub domain of the multimeric CaMKIIα, leading to oligomeric stabilization, proposed as the mechanistic basis for the reported neuroprotective action of such analogs ^2–4^. CaMKIIα holds a key role in synaptic plasticity, learning and memory ^5^, and mutations in CaMKII human genes have been uncovered to cause intellectual disability ^6–10^. Furthermore, after ischemic brain injury, aberrant calcium (Ca^2+^) levels trigger cell death involving excessive CaMKIIα activity and, indeed, CaMKII peptide inhibitors have been found to be neuroprotective post-injury ^11,12^. Whereas these compounds inhibit via kinase domain interaction, GHB analogs are selective hub ligands and, thus, potential drug candidates for post-insult neuroprotection by a yet undisclosed allosteric mechanism-of-action.

**Figure 1.**
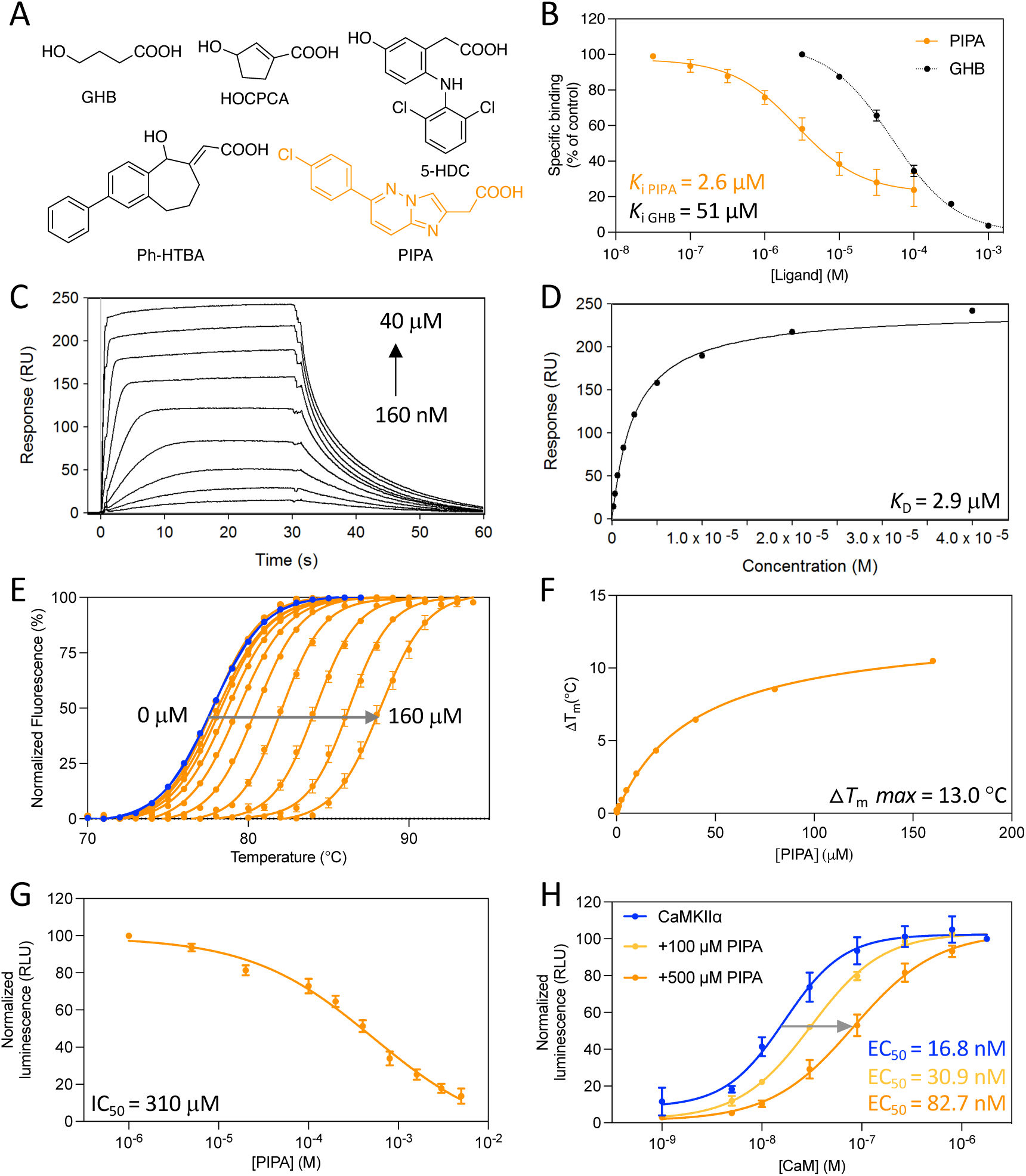
PIPA binds to CaMKIIα WT hub and inhibits substrate phosphorylation. **A)** Chemical structures of γ-hydroxybutyric acid (GHB), 3-hydroxycyclopenten-1-enecarboxylic acid (HOCPCA), 5-hydroxydiclofenac (5-HDC), (*E*)-2-(5-hydroxy-2-phenyl-5,7,8,9-tetrahydro-6*H*-benzo[7]annulen-6-ylidene)acetic acid (Ph-HTBA), and 2-(6-(4-chlorophenyl)imidazo[1,2-*b*]pyridazine-2-yl)acetic acid (PIPA). **B)** Concentration-dependent inhibition of [^3^H]HOCPCA binding by PIPA in whole-cell homogenates from HEK293T cells transfected with CaMKIIα (pooled data, n=4; GHB for comparison ^2^). **C)** Concentration-dependent binding of PIPA to immobilized CaMKIIα WT hub measured by SPR, and **D)** associated Langmuir-binding isotherm (representative data of n=3). **E)** Right-shifted thermal shift assay melting curves of CaMKIIα WT hub upon binding of PIPA (representative data of n=3). **F)** Saturation isotherm (representative). **G)** Concentration-dependent inhibition of CaMKIIα syntide-2 by PIPA in the luminescence-based ADP-Glo kinase assay (pooled data, n=3, mean ± SEM). **H)** Right-shifted CaM curve of CaMKIIα syntide-2 phosphorylation in the presence of PIPA (pooled data, n=3). For clarity, only mean values without variation are given; all mean values can be found in the respective results section.

CaMKIIα is a structurally and functionally complex protein. The holoenzyme is multimeric, where subunits are organized into predominantly dodecamers and tetradecamers (12 and 14 subunits). Each subunit consists of an *N*-terminal kinase domain, a regulatory segment, a linker region, and a *C*-terminal hub domain. The hub domain governs the oligomerization of individual subunits into functionally competent holoenzymes ^13^ and is also emerging as a regulator of kinase activity ^14^.

Although debated in the field ^15^, hub domain dynamics is also suggested to play a role in so-called activation-triggered subunit exchange which can lead to activation of neighboring holoenzymes ^16, 17^.

We recently revealed the first structural details of the small-molecule binding pocket of CaMKIIα hub from a 2.2 Å resolution co-crystal structure of the tetradecamer-stabilized human CaMKIIα hub mutant (CaMKIIα 6x hub ^18^) with the GHB analog 5-hydroxydiclofenac (5-HDC; Fig. 1A) ^2^. 5-HDC was found bound to 12 out of 14 subunits of the tetradecameric CaMKIIα hub oligomer, and to interact specifically with the positively charged residues Arg433, Arg453, and Arg469 as well as His395 that can be either neutral or positively charged. In addition, intrinsic tryptophan fluorescence (ITF) measurements with purified CaMKIIα hub domain demonstrated that binding of 5-HDC led to displacement of a flexible loop located at the edge of the binding pocket which contains the central residue tryptophan 403 (Trp403) ^2^.

The GHB analogs 3-hydroxycyclopent-1-enecarboxylic acid (HOCPCA; Fig. 1A) and (*E*)-2-(5-hydroxy-2-phenyl-5,7,8,9-tetrahydro-6*H*-benzo[7]annulen-6-ylidene)acetic acid (Ph-HTBA; Fig. 1A), known from competition studies to bind to the same site and increase oligomeric thermal stability. Especially, the affected oligomerization has been suggested to explain their neuroprotection by this mechanism ^3,19^. Recently, in biochemical studies Ph-HTBA ^19^, but not the smaller-size compound HOCPCA ^2^, was found to cause an outward flip of Trp403 located in the flexible hub domain loop, and to inhibit substrate phosphorylation of the CaMKIIα holoenzyme ^3^. Due to these observations, we hypothesize that distinct ligand-induced molecular interactions involving conformational changes of Trp403 and its CaMKIIα hub associated loop contribute further to the function of CaMKIIα hub ligands. In this study, we provide evidence for this hypothesis using model compounds binding competitively with GHB: the larger-type GHB analog 2-(6-(4-chlorophenyl)imidazo[1,2-*b*]pyridazine-2-yl)acetic acid) (PIPA), and the smaller size GHB analogs, acetate (Fig. 1A) ^20^ and HOCPCA. PIPA was previously reported as a high affinity GHB-site competitive ligand (*K*_i_ value of 0.22 μM in rat native synaptic membranes ^21^). Using X-ray crystallography, biophysical and biochemical approaches, we show that both PIPA and acetate bind to the CaMKIIα hub but produce distinct biochemical and structural outcomes. In particular, we confirm the effect of ligand size on hub domain loop movement and allosteric control of CaMKIIα function. Furthermore, we identify the small-molecule compound PIPA as a conformationally selective CaMKIIα hub ligand for studying new aspects of CaMKIIα hub conformational changes. Explicitly herein, we pinpoint Trp403 - unique to CaMKIIα - as a key molecular determinant.

## Results

### PIPA binds directly to the CaMKIIα hub domain

In extension to earlier reports on mid nanomolar affinity binding to the native GHB high-affinity site in rat cortical membranes ^2^, PIPA was verified to bind to recombinant CaMKIIα holoenzyme, albeit with micromolar affinity. This was shown by concentration-dependent inhibition of [^3^H]HOCPCA binding by PIPA to whole-cell homogenates of HEK293T cells transiently overexpressing homomeric CaMKIIα holoenzyme (*K*_i_ = 2.6 μM; p*K*_i_ = 5.65 ± 0.15) (Fig. 1B). To further confirm binding of PIPA directly to the CaMKIIα hub domain, surface plasmon resonance (SPR) binding studies were performed. The SPR data revealed binding of PIPA to immobilized CaMKIIα wild-type (WT) hub with a mean *K*_D_ value of 2.9 ± 0.6 μM (Fig. 1C-D). Similar binding affinities were found for PIPA to the CaMKIIα 6x hub construct and the CaMKIIα holoenzyme (Fig. S1). Thus, although affinity of PIPA is compared to native conditions, these experiments show that PIPA binds directly to the isolated hub domain.

### Thermal stabilization of the CaMKIIα hub domain by PIPA

An increased thermal stability of the purified CaMKIIα WT hub has been shown as a characteristic upon binding of GHB and several other known GHB analogs ^2, 22^. By using the described thermal shift assay based on differential scanning fluorometry, the *T*_m_ for the CaMKIIα WT hub was found to be 77.4 ± 0.2°C (Fig. 1E). Upon PIPA binding, a concentration-dependent increase in CaMKIIα WT hub stability was observed (Α*T*_m_ = 13.04 ± 0.09 °C) (Fig. 1E-F). However, the maximum Α*T*_m_ for the CaMKIIα WT hub upon PIPA binding was found to be 10.7 °C less compared to the analog 5-HDC which had a maximum Α*T*_m_ of 23.7 ± 0.2 °C (Fig. S2A-C). On the other hand, binding of the smaller-size compound HOCPCA induced a similar extent of hub stabilization as PIPA, with a maximum Α*T*_m_ of 11.5 ± 0.6 °C (Fig. S2D-F).

### PIPA inhibits CaMKIIα-mediated substrate phosphorylation

We recently reported that GHB and the two CaMKIIα hub ligands HOCPCA and Ph-HTBA mediate neuroprotection in mouse models of stroke ^2, 3^. Common to these ligands is their ability to bind to CaMKIIα and increase CaMKIIα hub thermal stability. For the larger-type analog Ph-HTBA, it was further seen that activity may also involve altered holoenzyme dynamics due to a reduced kinase activity ^3, 19^. Specifically, Ph-HTBA was found to inhibit CaMKIIα-dependent syntide-2 phosphorylation in the ADP-Glo kinase assay under sub-maximal calmodulin (CaM) concentration (30 nM) ^3^. Similarly, we observed that PIPA inhibited syntide-2 phosphorylation (IC_50_ = 388.6 μM; pIC_50_ = 3.44 ± 0.09) (Fig. 1G). A reduced CaM sensitivity was observed with increasing PIPA concentration with EC_50_ of CaM of 30.9 µM using 100 µM PIPA and 82.7 µM using 500 µM PIPA (Fig. 1H). Thus, to further compare, we also tested the smaller-size compounds HOCPCA and acetate (Fig. 1A), using corresponding sub-maximal (30 nM) CaM concentrations. Under this condition, neither of the compounds affected substrate phosphorylation, even at concentrations up to 5-10 mM (Fig. S3). Of note, in previous studies GHB analogs were all found to be inactive in the ADP-Glo kinase assay when performed at maximal CaM concentrations (1781 nM) ^2^.

### PIPA flips and restricts Trp403 positioning

Using the tetradecamer-stabilized CaMKIIα 6x hub ^18^, we obtained a co-crystal structure of the CaMKIIα hub oligomer bound to PIPA and acetate determined at 2.1 Å resolution (Fig. 2 and Table S1). In this structure, PIPA occupied two out of seven subunits in the equatorial plane, while the rest of the subunits were occupied by the low-affinity acetate (from the buffer) (Fig. 2A, B). In one of the subunits, PIPA bound at the conventional binding site, whereas in the other subunit it was located in the vicinity of the conventional binding site and with acetate and a polyethylene (PEG) moiety observed in the conventional binding site. Intriguingly, PIPA and acetate differentially directed movements of Trp403 in the loop at the edge of the binding pocket.

**Figure 2.**
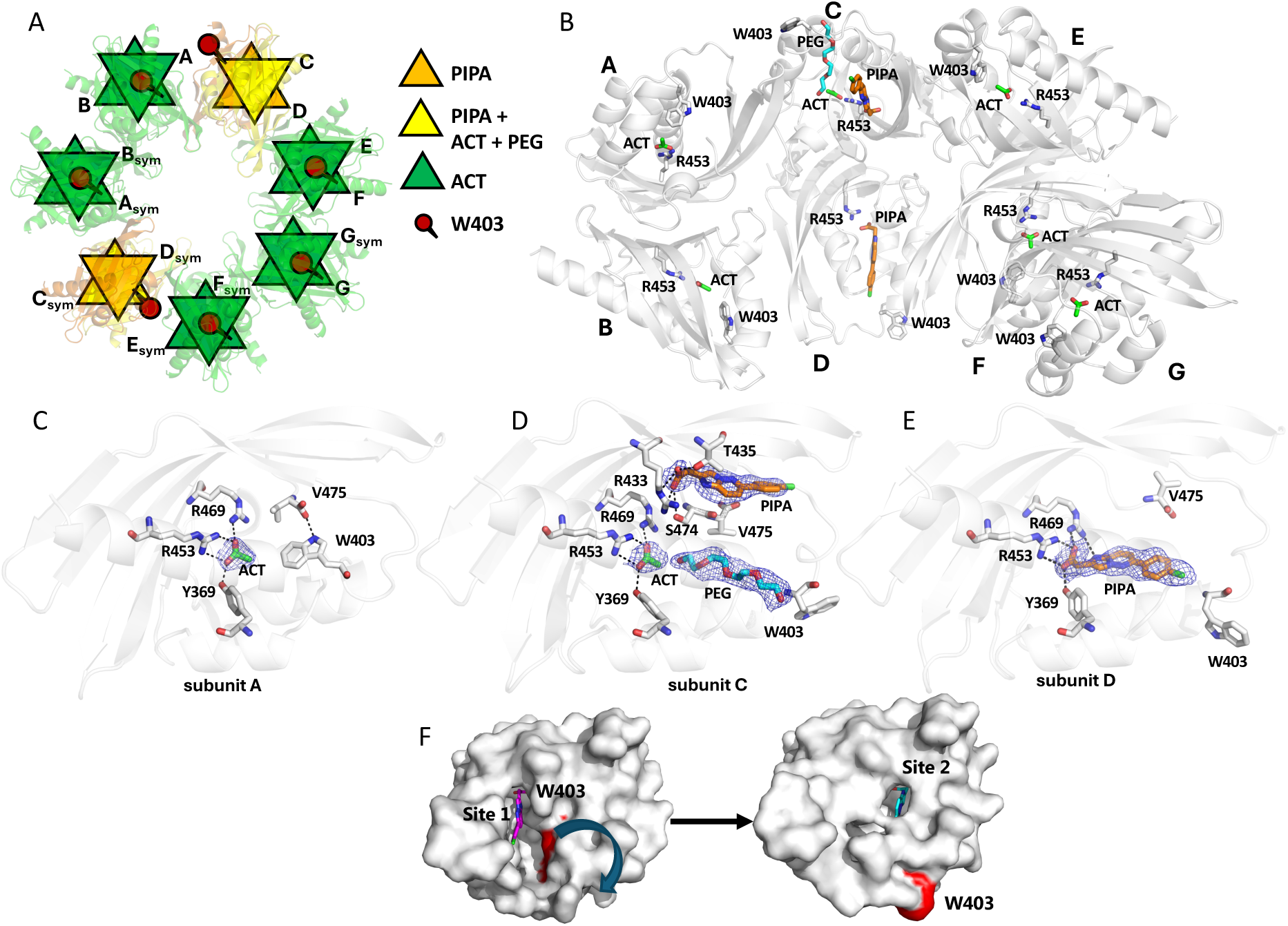
PIPA bound to CaMKIIα 6x hub promotes Trp403 flip while acetate does not. **A)** X-ray crystal structure of tetradecameric CaMKIIα 6x hub in schematic representation, with two subunits bound to PIPA (orange, yellow) and six subunits to acetate (ACT) only (green) or ACT+PIPA (yellow). Trp403 is shown in red. **B)** The structure of CaMKIIα 6x hub with seven subunits A-G in the asymmetric unit of the crystal, forming a tetradecamer with symmetry-related molecules, is shown in white cartoon representation with PIPA (orange carbon atoms), ACT (green), PEG (cyan), Arg453 (white), and Trp403 (white) shown in sticks representation. **C)** Zoom-in on binding mode and interactions of acetate in subunit A. Salt bridges and hydrogen bonds between acetate and corresponding amino-acid residues are shown as black dashed lines and with compounds and residues as sticks representation. In addition, a hydrogen bond between the C-terminal carboxylate group of Val475 and Trp403 is shown. The 2F_o_-F_c_ electron densities for the ligands are contoured at 1.0 sigma and carved at 1.6 Å. **D)** Alternative binding mode and interactions of PIPA in subunit C. Hydrogen-bonding interactions between PIPA, ACT, and corresponding amino-acid residues are shown as black dashed lines and with compounds and residues as sticks representation. The 2F_o_-F_c_ electron densities for PIPA (carved at 1.6 Å), ACT (carved at 1.75 Å), and PEG (carved at 2 Å) have been contoured at 0.5 sigma. In addition, Trp403 and Val475 are shown as sticks representation. **E)** Binding mode and interactions of PIPA in subunit D. Salt bridges and hydrogen-bonds between PIPA and corresponding amino-acid residues (as sticks representation) are shown as black dashed lines. In addition, Trp403 and Val475 are shown as sticks representation. The 2F_o_-F_c_ electron density for PIPA has been contoured at 0.5 sigma and carved at 1.8 Å. **F**) Surface representation of subunit A, where Trp403 (marked in red) is flipped inwards as shown to the left. PIPA (magenta sticks) has been modelled into site 1 to illustrate entrance of PIPA into the CaMKIIα 6x hub. Subunit D is shown to the right where PIPA (cyan sticks) binds at the conventional binding site 2, where Trp403 is flipped outwards as shown to the right.

The key molecular Interactions of the compounds (PIPA and acetate) are illustrated in Fig. 2. Acetate binds into the hub cavity of CaMKIIα 6x hub with the carboxylate moiety forming salt bridges with Arg453 and Arg469, and a charge-assisted hydrogen bond with Tyr369 (Fig. 2C, subunit A) as also observed for PIPA. The binding of the smaller-size acetate allows Trp403 to adopt an inward-flipped conformation and the C-terminal carboxylate group of Val475 to form a hydrogen bond to the indole nitrogen of Trp403 (Fig. 2A, B, C). In subunit C, PIPA occupies the binding pocket together with acetate and a PEG molecule from the crystallization buffer. Here, PIPA binds at a site (site 1) in the vicinity of the conventional binding site (site 2), with the carboxylate group of PIPA forming hydrogen-bonding interactions with the hydroxyl groups of Ser474 and Thr435 and a potential salt bridge with Arg433 (Fig. 2D). The PEG molecule binds at the conventional site near acetate (Fig 2D). In subunit C, Trp403 adopts an outward-facing conformation (Fig. 2A, B, D). In subunit D, PIPA was observed to bind to the conventional binding site (site 2) as previously observed for 5-HDC, with the carboxylate group of PIPA forming a salt bridge with Arg453 and Arg469, and a charge-assisted hydrogen bond to Tyr369 (Fig. 2E). In addition, a potential hydrogen bond is formed between the heteroaromatic bicyclic ring system of PIPA and Arg469. The binding of PIPA leads Trp403 to flip outward in a similar fashion as observed in subunit C (Fig. 2A, B, D, E).

Altogether, the structural data suggests that PIPA binds to the CaMKIIα 6x hub in a two-step process. First, PIPA enters the binding site via the accessible intermediate site 1 (represented by subunit A) and causes Trp403 and the connecting loop to flip outwards. Then PIPA moves into the more buried site 2 (represented by subunit D) where it binds to Arg453 and restricts the Trp403 in the outward-flipped position (Fig 2F).

### CaMKIIα WT hub self-association is induced by ligand binding

Dissociation from the dodecameric or tetradecameric CaMKIIα holoenzyme has been shown to be extremely slow with a *K*_D_ estimated at 10–20 nM ^23^. As the CaMKIIα holoenzyme concentration in dendritic spines is estimated to be 100 μM ^24^, it is therefore likely that these dodecameric or tetradecameric oligomers represent functionally relevant native structures of this protein. As the GHB analogs induce large stabilizing effects by binding to oligomers of the isolated CaMKIIα WT hub domain, these analogs were prior to this study hypothesized to change or stabilize the proportion of CaMKIIα hub dodecamers and tetradecamers, also in the CaMKIIα holoenzyme. Therefore, the structures of the CaMKIIα WT hub and the tetradecamer-stabilized CaMKIIα 6x hub mutant were investigated using small-angle X-ray scattering (SAXS), to elucidate the solution state and potential structural changes upon ligand binding. Furthermore, as Trp403, based on the X-ray crystallography data, seemed to play a central role in the binding of PIPA, we also investigated the ligand interaction with the CaMKIIα W403L hub mutant.

The SAXS data of neither CaMKIIα WT hub nor CaMKIIα 6x hub domains revealed any concentration-dependent effects within the concentration range examined (Fig. S4A-D). Comparing the three CaMKIIα hub domain constructs, distinct differences between the CaMKIIα WT hub and the tetradecamer-stabilized CaMKIIα 6x hub were observed, while the CaMKIIα W403L hub resembled the CaMKIIα WT hub (Fig. 3A). All three constructs exhibited the characteristics of folded multidomain structures as seen in the dimensionless Kratky plot (Fig. S4E). The CaMKIIα 6x hub appeared slightly larger both in the scattering curve (Fig. 3A) and upon indirect Fourier transformation to the pair distance distribution function (Fig. S4F) than the CaMKIIα WT hub and CaMKIIα W403L hub. Based on this observation and knowledge from the crystal structures and previous studies, we tested the respective fits of dodecamers and tetradecamers as well as mixtures of the two to the data. The best fits were obtained by the use of single oligomers, i.e, tetradecamers for the CaMKIIα 6x hub and dodecamers for the CaMKIIα WT hub (Fig. 3B) as also seen in the respective crystal structures (PDB entries 6OF8 and 5IG3) ^18, 25^. The dodecamer fit to the CaMKIIα WT hub is not perfect, and we cannot exclude e.g., minor fractions of tetradecamers or other aggregates (observed upon storage); however, the fit was not improved by inclusion of tetradecamers. Although some reports on the isolated CaMKIIα WT hub have suggested a 1:1 distribution of dodecamers and tetradecamers ^18, 25^, our data suggests dodecamers as the absolute prominent species for the CaMKIIα WT hub, consistent with observations on the CaMKIIα holoenzyme ^26^.

**Figure 3.**
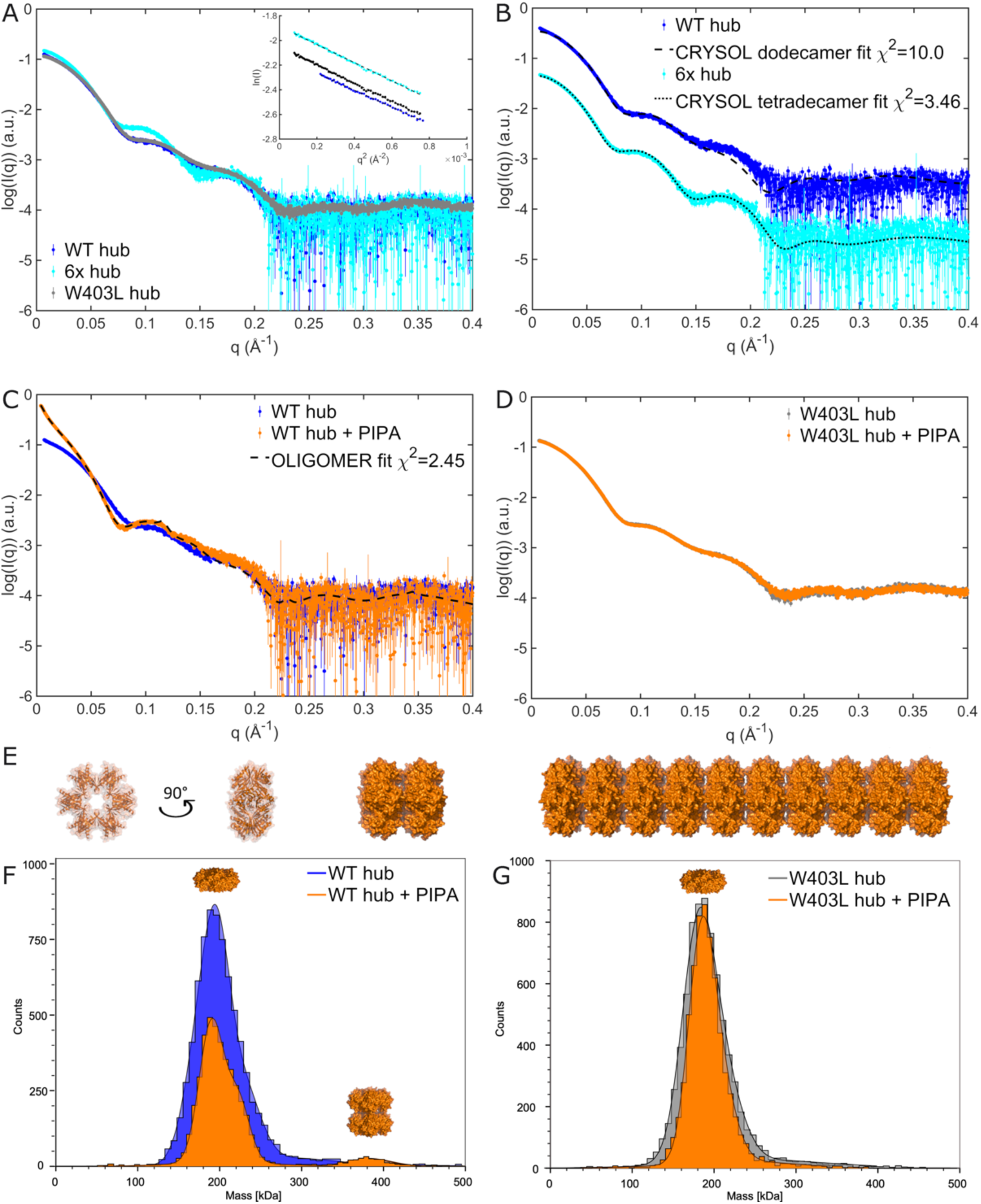
Solution structures of CaMKIIα WT hub reveals PIPA induced structured multimerization. **A)** SAXS scattering curves for the CaMKIIα WT, 6x, and W403L hub domains, with corresponding Guinier fits (insert). Full concentration series of CaMKIIα WT hub and CaMKIIα 6x hub are included in Fig. S4A-D. **B)** Comparison of CaMKIIα WT hub and CaMKIIα 6x hub curves to atomic models of the dodecamer (PDB entry 5IG3 ^25^) and tetradecamer (PDB entry 6OF8 ^18^), respectively. The scattering curves have been translated for clarity. **C)** SAXS scattering curves of CaMKIIα WT hub in the presence and absence of PIPA, including oligomer fit using model of self-associated dodecamers. **D)** SAXS scattering curves of CaMKIIα W403L hub in the presence and absence of PIPA. **E)** Examples of the stacked self-associated dodecamers used to model CaMKIIα WT hub with PIPA data. A single dodecamer is shown in two orientations. In addition, a dimer of dodecamers and a stacked model including 10 dodecamers are illustrated. Models and figure in E created using PyMOL ^46^. **F**) Representative MP histograms showing CaMKIIɑ WT hub diluted to 500 nM with and without the addition of PIPA. **G)** Representative MP histograms showing W403L CaMKIIɑ hub diluted to 500 nM, with and without PIPA. Each count indicates a single molecule. Additional MP data are shown in Fig. S6.

Whereas neither 5-HDC nor HOCPCA had any observable influence on the oligomeric state of the CaMKIIα WT hub construct (Fig. S4G-H), PIPA induced self-association of the CaMKIIα WT hub (Fig. 3C). A varying degree of self-association was observed in different preparations of CaMKIIα WT hub with PIPA, depending on the protein concentration upon buffer exchange and the storage time.

Based on the appearance of the scattering curves and the indication of a Bragg peak in one curve (at q = 0.115 Å^-1^, corresponding to a repeated real space distance of 54.5 Å), we hypothesized structured self-association of the oligomers through stacking as illustrated in Fig. 3E. A range of stacked dodecamer models were created and fitted to the scattering data of the CaMKIIα WT hub together with PIPA (Fig. 3C, Fig. S5 and Table S2). The highest concentration sample showed signs of aggregation immediately upon buffer exchange. SAXS data was in that case collected on the soluble fraction after removal of the non-soluble aggregates by centrifugation (Fig. 3C). This soluble fraction was fitted by a combination of dodecamers (∼37%), dimers of dodecamers (∼15%), and significant fractions of larger multimers (using models composed of 3-40 dodecamers; examples illustrated in Fig. 3E). Despite differences in the sample preparation, storage time, and data collection, convincingly, these stacked models could fit the different data measured with volume fractions of the different components consistent with the variation in concentration of the protein during sample preparation (Fig. S5 and Table S2). Although the finite size of these stacks may exceed the resolution limit of the data and the resulting fits do not exclude a broader variation, the trend from these fits consistently shows that PIPA under these experimental conditions induces formation of self-associated stacks. The self-association induced by PIPA was observed for the CaMKIIα WT hub domain in the entire concentration range examined (1.2 to 3.9 mg/mL), while no self-association was detected for the CaMKIIα 6x hub and CaMKIIα W403L hub constructs with PIPA despite investigating higher protein concentrations (6.7 and 6.8 mg/mL, respectively) (Fig. 3D and S4H).

To validate the occurrence of PIPA-induced CaMKIIα WT hub self-association at lower protein concentrations, mass photometry studies were performed. In the absence of PIPA, CaMKIIα WT hub and CaMKIIα W403L hub both adopted a structure with molecular weight (MW) around 188-198 kDa, compatible with dodecamers (Fig. 3F-G and Fig. S6). However, CaMKIIα WT hub with PIPA displayed a large shoulder on the main peak around 220 kDa as well as a much larger species at 372 kDa (corresponding to two dodecamers) (Fig. 3F and Fig. S6). CaMKII W403L hub did not display any change in molecular weight upon PIPA addition, indicating that there was no significant change in its oligomerization state. Based on SAXS and mass photometry data, we conclude that PIPA introduces ordered self-association of CaMKIIα WT hub and that this self-association involves Trp403 as mutation of this residue prevents formation of larger species.

### The Trp403Leu mutation increases PIPA-induced hub stability

As conformational change of the CaMKIIα-specific residue Trp403 at the upper edge of the binding pocket has previously been demonstrated for the compound 5-HDC using ITF measurements ^2^, this was also probed for PIPA. A concentration-dependent quenching of a fluorescence signal of CaMKIIα 6x hub in the presence of PIPA was observed (IC_50_ = 5.1 μM; pIC_50_ = 5.30 ± 0.06) (Fig. 4A-B). This further supported the observation of an outward flip of Trp403 introduced by PIPA in the X-ray structure of CaMKIIα 6x hub. These data supports that PIPA reaches up into the hydrophobic space in the upper cavity of the CaMKIIα hub domain and leads to an outward placement of Trp403.

**Figure 4.**
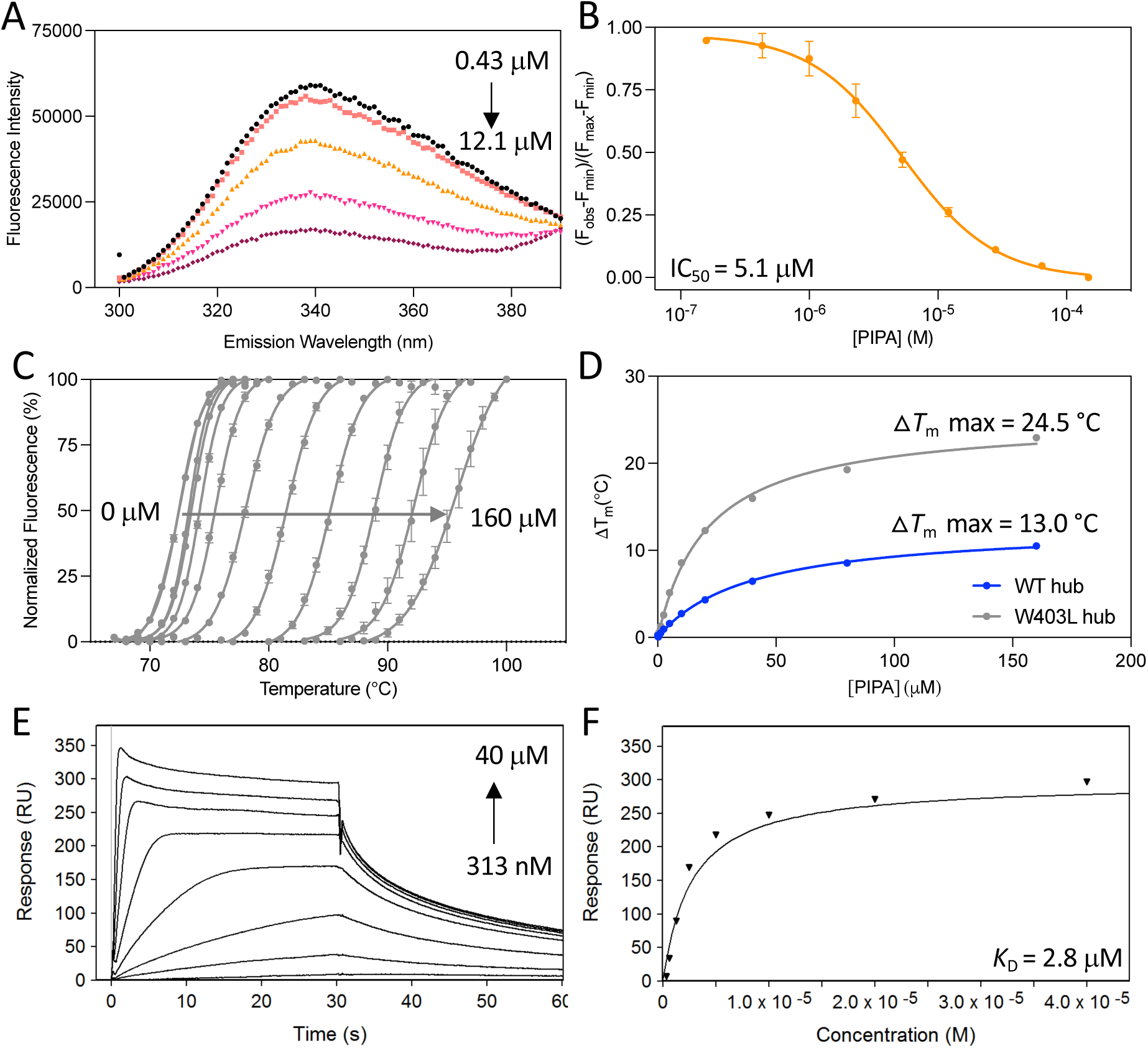
CaMKIIα hub mutation confirms a central role for Trp403 in PIPA binding. **A)** Quenching of intrinsic tryptophan fluorescence caused by Trp403 flip in the CaMKIIα 6x hub with increasing concentrations of PIPA, and **B)** resulting normalized inhibition curve (pooled data, n = 3). **C)** Right-shifted thermal shift assay melting curves of CaMKIIα W403L hub upon binding of PIPA (representative data). **D)** Thermal melting point of CaMKIIα WT hub (blue) and CaMKIIα W403L hub (grey) plotted against increasing concentrations of PIPA (representative data of n=3). **E)** Concentration-dependent binding of PIPA to immobilized CaMKIIα W403L hub measured by SPR (representative data), and **F)** associated Langmuir-binding isotherm (representative data of n=2).

To investigate the impact of the ligand-induced Trp403 conformational restriction on thermal stability, the CaMKIIα W403L hub was evaluated in the thermal shift assay in the presence of ligands. In the absence of ligand, the *T*_m_ for the CaMKIIα W403L hub was found to be 73.0 ± 0.4 °C, compared to 77.4 ± 0.2 °C for the CaMKIIα WT hub. Thus, replacing Trp403 with a leucine slightly reduced the thermal stability of the hub. However, in the presence of PIPA, the maximum Λ1*T*_m_ for the CaMKIIα W403L hub was found to be excessively increased (24.5 ± 0.4 °C) (Fig. 4C), compared to only 13.04 ± 0.09 °C for the CaMKIIα WT hub (Fig. 4D). In contrast, the maximum Λ1*T*_m_ for the CaMKIIα W403L hub in the presence of 5-HDC and HOCPCA was only slightly increased relative to the CaMKIIα WT hub with these ligands (from 11.5 ± 0.6 °C to 15.2 ± 1.0 °C with HOCPCA and from 24.5 ± 0.4 °C to 28.21 ± 0.03 with 5-HDC) (Fig. S2A-F). To explore potential changes in the binding affinity of PIPA to the CaMKIIα W403L hub domain compared to CaMKIIα WT hub, SPR binding studies were performed. The SPR data revealed unchanged binding between PIPA and immobilized CaMKIIα W403L hub (*K*_D_ of 2.80 ± 0.13 μM) (Fig. 4E-F) and CaMKIIα WT hub.

## Discussion

The CaMKIIα hub domain is proposed to be involved in allosteric regulation of the kinase activity of the CaMKIIα holoenzyme ^14^. This study adds experimental support to this notion and suggests that this is governed by the placement of the highly flexible loop containing Trp403. The GHB analog PIPA was found to specifically bind to the CaMKIIα hub domain, and albeit with low potency, to inhibit the CaMKIIα holoenzyme kinase activity. Due to the requirement of sub-maximal CaM concentrations to observe an effect of PIPA, and the apparent high micromolar potency of PIPA to achieve kinase inhibition compared to the high nanomolar hub affinity, it cannot be excluded that PIPA interacts with other sites at CaMKIIα (awaits further studies). However, SPR data on the CaMKIIα holoenzyme compared to CaMKIIα WT hub shows comparable binding and suggests specific interactions of PIPA to the hub domain. As such, PIPA together with Ph-HTBA^3^ represent first-in-class allosteric inhibitors of CaMKIIα enzymatic activity acting through binding to the CaMKIIα hub domain. Furthermore, in the isolated CaMKIIα 6x hub domain PIPA led to a conformational restriction of the loop containing Trp403 flipped outwards. We further observed that the much smaller compound acetate binds to the same pocket and binding site residues as PIPA. However, with acetate bound, the Trp403 is pointing inwards as also observed for CaMKIIα 6x hub without ligand (PDB entry 6OF8). Thus, the similarly small compounds GHB and HOCPCA binding within the same CaMKIIα hub pocket, may likewise be predicted to allow an inward positioning of Trp403, possibly stabilizing a slightly different conformational state of the CaMKIIα hub domain with yet unknown biological significance.

Evidently, no Trp403 flip was observed for either of these two compounds based on ITF measurements, and no kinase inhibition could be demonstrated ^2^. Thus, it appears that there is a correlation between the extent of the displacement of the flexible loop containing Trp403 and substrate phosphorylation. In other words, minute differences in the binding mode of these compounds seem to dictate functional outcomes. Such differences are seemingly linked to the molecular size of the ligands where PIPA, because of a larger size, sterically prevents an inward-posing of the loop containing Trp403 and restricts it in an outward-flipped conformation.

We further revealed that PIPA stabilizes the CaMKIIα WT hub to a significantly lesser extent than 5-HDC, shown by a 10.7 °C lower maximum Δ*T*_m_ in the thermal shift assay. However, upon binding to the CaMKIIα W403L hub, both compounds were equally effective at stabilizing the domain. Overall, this suggests that an outward-flipped Trp403, provoked by PIPA, results in a lesser stabilization of the CaMKIIα hub-PIPA complex compared to when Trp403 is not present (CaMKIIα W403L hub).

Moreover, it suggests that the Trp403 is not similarly restricted when binding to 5-HDC, although previously shown to cause a Trp flip ^2^. This was also supported when comparing the structure of CaMKIIα 6x hub with PIPA and the previously published structure with 5-HDC ^2^, which revealed subtle differences in the extent of Trp403 flipping induced by the two ligands (Fig. S7). In the structure with 5-HDC, more space is observed potentially allowing for Trp403 movement, while the steric bulk of PIPA appears to push and restrict the Trp403 loop into an even more outward pose than 5-HDC. Overall, the data suggests that conformational changes of Trp403 and its connecting loop is a determinant for oligomeric stability and, indirectly, functional activity.

As the GHB analogs induce large stabilizing effects by binding to the CaMKIIα hub domain, they were prior to this study hypothesized to change or stabilize the proportional distribution of the different CaMKIIα hub oligomers. Importantly, the SAXS data in this study did not support any change in the proportions of dodecameric and tetradecameric forms of CaMKIIα WT hub upon ligand binding.

However, the data do not exclude the possibility that these ligands bind and stabilize the existing complexes of the hub domains, and thus prevent any natural dynamic exchange between these, e.g. release and exchange of subunits ^27, 28^. Excitingly, the SAXS data revealed another unexpected effect of PIPA binding. For the CaMKIIα WT hub, PIPA induced highly structured self-association (dodecamer stacking as illustrated in Fig 3E). This effect was, however, not observed for the CaMKIIα WT hub alone nor for the CaMKIIα W403L hub. Furthermore, self-association of the CaMKIIα WT hub was not observed in the presence of HOCPCA or 5-HDC. Overall, this infers a probe-dependent effect on stacking involving the altered peripheral surface of the CaMKIIα WT hub oligomers when the Trp403-containing loop is restricted in an outward conformation by the bulky PIPA. A limitation in the SAXS study is the protein concentrations required for the measurements. Importantly, mass photometry data showed the appearance of PIPA-induced self-association down to 100 nM concentration of CaMKIIα WT hub. This observation further indicates that this structural effect could be physiologically important. Thus, it is tempting to speculate that the binding of PIPA promotes self-association of CaMKIIα WT hub by restricting Trp403 in the outward flip.

Interestingly, we did not observe PIPA induced stacking of CaMKIIα 6x hub under similar conditions as for CaMKIIα WT hub. This indicated that additional factors than Trp403 may influence the stacking, including the oligomeric state and the multiple mutations in CaMKIIα 6x hub. However, in the X-ray crystal packing of the CaMKIIα 6x hub with PIPA, the tetradecameric hub was found to be surrounded by tetradecamers positioned perpendicularly to this (face-to edge) as shown in Fig. S8. Although being different than the pattern of self-association observed in the CaMKIIα WT hub by SAXS, it is apparent that Trp403 plays an important role in self-association per se, as in each interface an outward-posing Trp403 in one tetradecamer participates in a π–π interaction to another outward-posing Trp403 from a connecting tetradecamer (Fig. S8). Investigating the previously published X-ray crystal structure of the CaMKIIα 6x hub without ligand (PDB entry 6OF8), a similar stacking pattern as observed in SAXS could be observed in the structure with a repetitive distance of 56.2 Å ^18^. This could suggest self-association as a general feature of the CaMKIIα hub domain, possibly depending on loop flexibility and the proportion of outward-posing Trp403 available for π– π-stacking.

Although binding of PIPA to the CaMKIIα holoenzyme inhibits substrate phosphorylation, it remains to be shown whether the observed self-association also happens to the CaMKIIα holoenzyme and how such potential stacking might affect holoenzyme activity under different physiological or pathophysiological conditions. Interestingly, activation-triggered self-association has previously been observed for the CaMKIIα holoenzyme and suggested as an inactivation mechanism of CaMKIIα following ischemia ^29–31^, which supports self-association as a potential relevant regulatory or protective mechanism. Altogether the presented data shed new light on allosteric regulation of CaMKIIα activity via the hub domain, where positioning of Trp403 in the flexible hub loop is central. Further mechanistic insight into this regulation is warranted in future experiments.

## Materials and Methods

### Cell culturing and transfection of CaMKIIα in HEK293T cells

Human embryonic kidney (HEK) 293T (#CRL-3216, ATCC) cells were maintained in DMEM GlutaMAX medium (#61965026, Gibco) supplemented with 10% fetal bovine serum and 1% penicillin-streptomycin (#15140122, Invitrogen), in a humidified 5% CO_2_ atmosphere at 37°C. HEK293T cells were transfected with rat CaMKIIα WT (pCMV6-CaMKIIα-Myc-DDK, #RR201121, Origene), which was generated and sequence-verified by GenScript Biotech (Leiden, the Netherlands). Transfections were performed using polyethyleneimine (PEI) (Polysciences Inc., Warrington, PA, USA). The day before transfection, cells were seeded in 10 cm culture dishes at a density of 2.0 x 10^6^. On the day of transfection, 8 μg plasmid DNA was diluted in 970 µL serum-free medium and 24 μL 1 mg/mL PEI added. After 15 min of incubation at room temperature, the DNA/PEI mixture was added to the cells. Transfected cells were used for generating lysates for whole-cell homogenate binding, as described below.

### [^3^H]HOCPCA binding assay to whole cell homogenates

This assay was performed as previously described ^2^. The binding experiments were performed in 48-well format using glass tubes (#10682424, Corning™) with 40 nM [^3^H]HOCPCA, 100 μg HEK293T whole cell homogenate and test compound in a total volume of 400 μL binding buffer (50 mM KH_2_PO_4_, pH 6.0). GHB (3 mM) was used for determining non-specific binding. Binding equilibrium was achieved by 1 hr incubation at 0-4 °C, followed by protein precipitation with addition of 1.6 mL acetone for 1 hr at -20 °C. Protein was collected on GF/C unifilters by rapid filtration with ice cold binding buffer using a Brandel M48-T cell harvester (Alpha Biotech). Finally, filters were submerged in 3 mL OptiFluor scintillation liquid (Perkin Elmer) to allow measurement of radioactivity (DPM) for 3 min per sample. Experiments were performed in technical triplicates and curves generated from pooled data from four individual experiments. *K*_i_ values were calculated from the obtained IC_50_ values using the Cheng-Prusoff equation and previously determined *K*_D_ value ^2^. Mean p*K*_i_ values are reported as mean ± SEM.

### Expression and purification of CaMKIIα hub domains

The human CaMKIIα WT hub (UniprotKB Q9UQM7, residues 345-475), CaMKIIα W403L hub and CaMKIIα 6x hub (containing the six mutations Thr354Asn, Glu355Gln, Thr412Asn, Ile414Met, Ile464His, and Phe467Met) were expressed and purified as previously described ^2, 18, 32^. In brief, recombinant CaMKIIα hub domains with an *N*-terminal 6x His-precision protease expression tag were inserted into pSKB2 (6x hub) and pET28a (WT hub and W403L hub) vectors with kanamycin resistance, respectively. The proteins were expressed in BL21 (DE3) *E. coli* cells and purified by His-tag protein capture to nickel IMAC columns (HisTrap FF) and eluted using 75% 25 mM Tris, 150 mM KCl, 1M imidazole, 10% glycerol, pH 8.5. The proteins were buffer exchanged into a buffer containing 25 mM Tris, 150 mM KCl, 10 mM imidazole, 1 mM DTT, 10% glycerol, pH 8.5 using a HiPrep 26/10 column before precision protease was added overnight to remove the 6x His-tag. Lastly, the proteins were further purified using reverse HisTrap and size exclusion chromatography (Superose-6 gel filtration column). All purification steps were performed at 4 °C and all columns were purchased from Cytiva.

### Surface plasmon resonance

SPR measurements were performed at 25 °C using a Pioneer FE instrument (Sartorius). Recombinant CaMKIIα WT hub, CaMKIIα 6x hub, CaMKIIα W403L hub, and CaMKIIα holoenzyme (#02-109, Carna Biosciences) were immobilized by amine coupling on to a biosensor using 20 mM NaAc, pH 5 immobilization buffer. PIPA was injected in 2-fold concentration series over the immobilized CaMKIIα hub constructs using a MES running buffer (20 mM MES, 150 mM NaCl, 1 mM DTT, pH 6) and CaMKIIα holoenzyme using an HBS-P running buffer (10 mM Hepes, 150 mM NaCl, 0.005% tween, 1 mM DTT, pH 7.4). The data was analyzed using Qdat Data Analysis Tool version 2.6.3.0 (Sartorius). The sensorgrams were corrected for buffer bulk effects and unspecific binding of the samples to the chip matrix by blank and reference surface subtraction (activated flow cell channel by injection of EDC/NHS and inactivated by injection of ethanolamine). The dissociation constants (*K*_D_) were estimated by plotting responses at equilibrium (R_eq_) against the injected concentration and curve fitted to a Langmuir (1:1) binding isotherm. The calculated *K*_D_ value is mean ± SEM of three individual experiments.

### Thermal shift assay

Thermal melting points of CaMKIIα WT hub was determined without compound and in the presence of 0.3 - 160 µM PIPA by differential scanning fluorimetry (DSF). Each sample was prepared with a final concentration of 0.1 mg/mL CaMKIIα and 8x SYPRO® Orange Protein Gel Stain (Life Technologies, #S6650) in MES buffer (20 mM MES, 150 mM NaCl, 1 mM DTT, pH 6) to a volume of 25 µL/well in a 96-well qPCR plate. Fluorescence was measured on a Mx3005P qPCR system (Agilent Technologies) using 492 nm as excitation and 610 nm as emission in 85 cycles with a 1 °C temperature increase; 25-100 °C. Data analysis was performed in GraphPad Prism (v. 10) where the sigmoidal curves of normalized fluorescence intensity was fitted to the Boltzmann equation and *T*_m_ values were acquired. Further, the maximum difference in *T*_m_ (Δ*T*_m_ max) was found via non-linear regression (One site-Fit logIC_50_) from Δ*T*_m_ of each compound concentration compared to CaMKIIα WT hub plotted against compound concentration. Data (mean Δ*T*_m_ max values ± SEM) was obtained from three independent experiments performed in technical triplicates.

### ADP-Glo kinase assay

CaMKIIα activity was assessed using the ADP-Glo Kinase Assay kit (#V9101, Promega) with CaMKIIα holoenzyme (#PR4586C, Thermo Fisher). Kinase detection reagent and the ADP-Glo kinase reaction buffer (40 mM Tris, 0.5 mM CaCl_2_, 20 mM MgCl_2_, 0.1 mg/mL BSA, 50 μM DTT, pH 7.5) was prepared according to the manufacturer’s protocol. All experiments were performed with a working volume of 20 μL in 384-well white polypropylene plates (#784075, Greiner). The kinase reaction was performed with a final concentration of 3 ng CaMKIIα, 25 μM ATP, 50 μM Syntide-II, and varying compound and CaM concentrations. Inhibition curves were generated with 30 nM CaM and 1 – 5000 μM compound. CaM curves with 1 – 1781 nM CaM were obtained in the absence or presence of 100 μM or 500 μM compound. The kinase reactions with the abovementioned components were carried out for 55 min at 37 °C. Hereafter, excess ATP was depleted by incubation with 5 μL ADP-Glo Reagent for 40 min at room temperature. Finally, 10 μL Kinase Detection Reagent was added to each well to convert ADP to ATP and allow measurement after 30 min incubation at room temperature. Luminescence was measured on a LUMIStar Omega plate reader, and data analysis and curve fitting were performed using GraphPad (v. 9). IC_50_ and EC_50_ values were determined using ‘log(inhibitor) vs. response with variable slope’. Individual experiments were performed in technical triplicates and all curves are pooled data (mean ± SEM) of three independent experiments.

### Co-crystallization of PIPA with CaMKIIα 6x hub

Crystals were grown via sitting drop vapor diffusion at 20 °C with reservoir solution containing 11% w/v PEG3350, 300 mM potassium acetate, pH 8.0 as previously established for CaMKIIα hub crystals ^18, 25^. PIPA was co-crystallized by preincubating the protein solution (16 mg/mL) with 2.65 mM PIPA for 1 hr at 20 °C. 1.5 μL sitting drops were dispensed by adding 500 nL of protein stock to 1000 nL of reservoir solution. Drops were equilibrated against 50 μL of reservoir solution. Crystals appeared in 2 weeks and were cryoprotected (11% PEG, 300mM K acetate pH 8.0, 25% glycerol and 2.65mM PIPA) prior to flash cooling in liquid nitrogen for data collection. For statistics on X-ray diffraction data see Table S1.

### X-ray structure determination

X-ray diffraction data was collected at the Advanced Light Source (ALS) beamline 8.2.2 at wavelength 1.0000 Å and temperature of 100 K. Data was processed with XDS ^33^, scaled and merged with Aimless ^34^ in the CCP4 suite ^35^). The space group was *C*222_1_ with cell dimensions a = 103.05 Å, b = 182.92 Å, c = 107.76 Å, α = β = ψ = 90°. Initial phases were obtained by molecular replacement using Phenix Phaser-MR ^36^ and the structure of CaMKIIα 6x hub with 5-HDC (PDB entry 7REC ^2^) as search model. AutoBuild was next used for model building, refinement, and density modification. Further refinement of the structure was performed using Phenix (version 13) ^37^ and Coot ^38^ was used for manual model building. The CIF dictionary file for PIPA was generated using Phenix eLBOW ^39^. The 2F_o_-F_c_ electron density for PIPA and PEG is shown in Fig. 2. Statistics for structure refinement are available in Table S1.

### Small-angle X-ray scattering

For the SAXS studies CaMKIIα hub proteins were buffer exchanged into a MES buffer (20 mM MES, 150 mM NaCl, 1 mM DTT, pH 6.0) without or with the presence of ligands (PIPA 200 μM, HOCPCA 1000 μM, 5-HDC 100 μM) using Zeba™ Spin Desalting Columns (Thermo Fisher Scientific, 0.5 ml 7K MWCO). Dilution series were made of CaMKIIα WT hub, CaMKIIα 6x hub and CaMKIIα W403L hub in concentrations ranging from 1 to 7 mg/mL. Several preparations were made of CaMKIIα WT hub with PIPA at different protein concentrations prior to buffer exchange.

During the preparation of CaMKIIα WT hub with PIPA (sample iv) significant aggregation was visible immediately upon buffer exchange; thus, the sample was centrifuged (2 min, 4 °C, 10.000 g) and SAXS data was measured only on the soluble fraction (supernatant).

SAXS data was collected at the CPHSAXS facility (University of Copenhagen, Denmark) and at the P12 beamline operated by EMBL Hamburg at the PETRA III storage ring (DESY, Hamburg, Germany) ^40^. Data from the CPHSAXS facility was collected on a BioXolver L (Xenocs) using metal jet source (Excillum) equipped with a Pilatus3 R 300K detector (Dectris). Samples were automatically loaded using the BioCUBE sample handling robot from a 96-well tray. Initial data processing was done using BioXTAS RAW ^41^. Data from P12 was collected on a Pilatus6M detector (Dectris). Samples were loaded automatically using the ARINAX BioSAXS sample changer and sample flow during exposure. Data reduction was done using the SASflow pipeline ^42^.

Scattering intensities were measured at room temperature (20-22 °C) as a function of the momentum transfer q = (4πsin2θ)/λ with 2θ being the scattering angle and λ the X-ray wavelength. Details of the individual measurements (sample, concentration, facility, wavelength, detector distances, and exposure times) are listed in Table S3.

Images were radially averaged and overlap of the individual frames checked before averaging. For two of the samples of CaMKIIα WT hub with PIPA (samples ii and iii) some heterogeneity (possible development or sedimentation) was observed, while the data was still averaged for further analyses. Corresponding buffer measurements were subtracted and, when relevant, the data from different configurations were merged and scaled by concentration to the final data files. Subsequent primary data analysis, Guinier analyses, indirect Fourier transformation, and molecular weight estimation was done using PRIMUS ^43^. CRYSOL ^44^ was used for the evaluation of single structures against the experimental data, and for mixtures OLIGOMER ^45^ was used to determine best fit and corresponding volume fractions.

The following high-resolution models were used for comparison and analysis: dodecamer (PDB entry 5IG3)^25^ and tetradecamer (PDB entry 6OF8) ^18^. In addition, models of stacked dodecamers and tetradecamer were made by simple translation of the individual structures using PyMOL ^46^.

### Mass photometry

Mass photometry experiments were performed on a Refeyn One MP. Standard calibration was performed using 2.25 nM apoferritin, 15 nM bovine serum albumin, and 15 nM thioglycolic acid with known molecular weights: 440 kDa, 66.5 kDa, and 92.11 kDa, respectively. 3 μL of standard protein was diluted with 17 μL of filtered buffer (25 mM Tris-HCL, 150 mM KCl, pH 8.0) and measured for 60 sec.

For experiments without PIPA, CaMKIIα WT hub and CaMKIIα W403L hub were diluted to final concentrations of 500 nM, 200 nM, and 100 nM in MES buffer (20 mM MES, 150 mM NaCl, 1 mM DTT, pH 6.0) and measurements were made for 60 sec. For experiments with PIPA added, 1.5 mg/mL CaMKIIα WT hub and CaMKIIα W403L hub were incubated with 200 μM PIPA for 90 min and desalted into MES buffer + PIPA (ZebaTM spin, 7K MWCO). The eluant was diluted to 500 nM, 200 nM, and 100 nM for each measurement. Buffer alone was measured before each measurement and histograms were generated using the Refeyn’s DiscoverMP software.

### Intrinsic tryptophan fluorescence (Trp flip) assay

Intrinsic tryptophan fluorescence (ITF) measurements targeting the Trp403 was performed as previously described^2^. In brief, CaMKIIα 6x hub protein and PIPA were diluted in assay buffer (10 mM HEPES, 150 mM NaCl, 1 mM DTT, pH 7.4) and mixed in a microplate to obtain a protein concentration of 5 μM CaMKIIα 6x hub and 3.4 μM CaMKIIα WT hub. For absorbance and background fluorescence measurements, compounds were mixed with buffer for each compound concentration. PIPA showed no fluorescent properties. All measurements were performed in black half-area 96-well format low-binding OptiPlates (#6052260, PerkinElmer) for fluorescence and half-area UV-Star microplates (#675801, Greiner Bio-One) for absorbance. All measurements were recorded at 25 °C on a Safire2 plate reader (Tecan). Emission was recorded in the wavelength range of 300-450 nm with 1 nm increments and an excitation wavelength of 290 nm with 5 nm bandwidths. Fluorescence intensities at 340 nm were used for data analysis. To check for inner filter correction, the absorbance was measured in the range of 270-400 nm. PIPA showed absorbance for the higher concentrations and was corrected for inner filter effect with a factor between 0.99-1.4 calculated from: (*F_obs_* - *B*) * 10^0.5 *h*(*A_ex_*+*A_em_*)^

The fluorescence intensities were normalized according to: 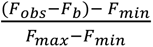

F_obs_ is the observed fluorescence intensity and F_b_ is the background fluorescence for compound in buffer alone. F_max_ is the fluorescence intensity of CaMKIIα hub alone without compound, and F_min_ is the fluorescence intensity when plateau is reached at high compound concentrations in the presence of the CaMKIIα hub domain. Since PIPA did not reach a plateau at high compound concentrations, F_min_ was set to the fluorescence intensity of buffer for all compounds tested.

Fluorescence intensities usually spanned from 3,000-40,000 for PIPA, while the fluorescence intensity for buffer was around 1,000. Non-linear regression was used for curve-fitting using the equation for ‘log(inhibitor) vs. response with variable slope’ to determine IC_50_ values (GraphPad Prism, v. 8).

## Author Contributions

D.N.: Methodology, Investigation, Formal analysis, Validation, Writing – Original Draft. A.S.G.L.: Investigation, Formal analysis. S.J.G.: Methodology, Investigation, Formal analysis, Validation, Writing – Review & Editing. R.A.: Investigation, Formal analysis. R.B.H: Investigation, Formal analysis. L.H.: Investigation, Formal analysis. J.B-J.: Investigation, Formal analysis. N.G-K.: Investigation, Formal analysis. C.L.G.: Methodology, Investigation, Formal analysis, Validation. B.F.: Validation, Supervision. M.M.S.: Methodology, Investigation, Formal analysis, Validation, Supervision, Writing. J.K.: Supervision. J.S.K.: Methodology, Investigation, Formal analysis, Validation, Supervision, Writing – Review & Editing. A.E.L.: Methodology, Investigation, Formal analysis, Validation, Writing – Review & Editing. P.W.: Conceptualization, Methodology, Formal analysis, Validation, Supervision, Writing – Review & Editing. S.M.Ø.S.: Conceptualization, Methodology, Investigation, Formal analysis, Validation, Supervision, Writing – Review & Editing.

## Data Availability

The structure coordinates and corresponding structure factor file of the CaMKIIα 6x hub in complex with PIPA, acetate, and PEG have been deposited in the Protein Data Bank under the accession code 9EOY. SAXS data and selected models are deposited in SASBDB^47^ under accession codes SASDUQ4, SASDUR4, SASDUS4, SASDUT4, SASDUU4, SASDUV4, SASDUW4, SASDUX4, SASDUY4, SASDUZ4, SASDU25, SASDU35, SASDU45, SASDU55, SASDU65, SASDU75.

## Acknowledgments

This work was supported through financial support from the following foundations and grants: The Lundbeck Foundation (R277-2018-260 to P.W.), The Novo Nordisk Foundation (NNF17OC0028664 and NNF21OC0067835 to P.W.), the Independent Research Fund Denmark (1026-00335B to P.W.), and the Drug Research Academy for Lundbeck Foundation pregraduate scholarships in pharmaceutical neuroscience (to A.S.G.L. and R.B.H.). We acknowledge the University of Copenhagen Small angle X-ray facility, CPHSAXS, funded by the Novo Nordisk Foundation (grant no. NNF19OC0055857) and assistance from Pernille S. Tuelung. https://drug.ku.dk/core-facilities/cphsaxs/. The synchrotron SAXS data was collected at beamline P12 operated by EMBL Hamburg at the PETRA III storage ring (DESY, Hamburg, Germany). We thank Cy Jeffries for the assistance in using the beamline as well as financial support through DANSCATT (funded by the Danish Agency for Science, Technology, and Innovation). X-ray diffraction data was collected at the Advanced Light Source beamline 8.2.2. The Berkeley Center for Structural Biology is supported in part by the Howard Hughes Medical Institute. The Advanced Light Source is a Department of Energy Office of Science User Facility under Contract No. DE-AC02-05CH11231. The ALS-ENABLE beamlines are supported in part by the National Institutes of Health, National Institute of General Medical Sciences, grant P30 GM124169.

## References

1. T. Bay, L. F. Eghorn, A. B. Klein and P. Wellendorph, GHB receptor targets in the CNS: focus on high-affinity binding sites, Biochem Pharmacol, 2014, 87, 220–228.

2. U. Leurs, A. B. Klein, E. D. McSpadden, N. Griem-Krey, S. M. Ø. Solbak, J. Houlton, I. S. Villumsen, S. B. Vogensen, L. Hamborg, S. J. Gauger, L. B. Palmelund, A. S. G. Larsen, M. A. Shehata, C. D. Kelstrup, J. V. Olsen, A. Bach, R. O. Burnie, D. S. Kerr, E. K. Gowing, S. M. W. Teurlings, C. C. Chi, C. L. Gee, B. Frolund, B. R. Kornum, G. M. van Woerden, R. P. Clausen, J. Kuriyan, A. N. Clarkson and P. Wellendorph, GHB analogs confer neuroprotection through specific interaction with the CaMKIIα hub domain, Proc Natl Acad Sci U S A, 2021, 118.

3. N. Griem-Krey, S. J. Gauger, E. K. Gowing, L. Thiesen, B. Frolund, A. N. Clarkson and P. Wellendorph, The CaMKIIα hub ligand Ph-HTBA promotes neuroprotection after focal ischemic stroke by a distinct molecular interaction, Biomed Pharmacother, 2022, 156, 113895.

4. N. Griem-Krey, A. B. Klein, B. H. Clausen, M. R. Namini, P. V. Nielsen, M. Bhuiyan, R. Y. Nagaraja, T. M. De Silva, C. G. Sobey, H. C. Cheng, C. Orset, D. Vivien, K. L. Lambertsen, A. N. Clarkson and P. Wellendorph, The GHB analogue HOCPCA improves deficits in cognition and sensorimotor function after MCAO via CaMKIIα, J Cereb Blood Flow Metab, 2023, 43, 1419–1434.

5. K. U. Bayer and H. Schulman, CaM Kinase: Still Inspiring at 40, Neuron, 2019, 103, 380–394.

6. P. H. Chia, F. L. Zhong, S. Niwa, C. Bonnard, K. H. Utami, R. Zeng, H. Lee, A. Eskin, S. F. Nelson, W. H. Xie, S. Al-Tawalbeh, M. El-Khateeb, M. Shboul, M. A. Pouladi, M. Al-Raqad and B. Reversade, A homozygous loss-of-function CAMK2A mutation causes growth delay, frequent seizures and severe intellectual disability, Elife, 2018, 7.

7. S. Kury, G. M. van Woerden, T. Besnard, M. Proietti Onori, X. Latypova, M. C. Towne, M. T. Cho, T. E. Prescott, M. A. Ploeg, S. Sanders, H. A. F. Stessman, A. Pujol, B. Distel, L. A. Robak, J. A. Bernstein, A. S. Denomme-Pichon, G. Lesca, E. A. Sellars, J. Berg, W. Carre, O. L. Busk, B. W. M. van Bon, J. L. Waugh, M. Deardorff, G. E. Hoganson, K. B. Bosanko, D. S. Johnson, T. Dabir, O. L. Holla, A. Sarkar, K. Tveten, J. de Bellescize, G. J. Braathen, P. A. Terhal, D. K. Grange, A. van Haeringen, C. Lam, G. Mirzaa, J. Burton, E. J. Bhoj, J. Douglas, A. B. Santani, A. I. Nesbitt, K. L. Helbig, M. V. Andrews, A. Begtrup, S. Tang, K. L. I. van Gassen, J. Juusola, K. Foss, G. M. Enns, U. Moog, K. Hinderhofer, N. Paramasivam, S. Lincoln, B. H. Kusako, P. Lindenbaum, E. Charpentier, C. B. Nowak, E. Cherot, T. Simonet, C. A. L. Ruivenkamp, S. Hahn, C. A. Brownstein, F. Xia, S. Schmitt, W. Deb, D. Bonneau, M. Nizon, D. Quinquis, J. Chelly, G. Rudolf, D. Sanlaville, P. Parent, B. Gilbert-Dussardier, A. Toutain, V. R. Sutton, J. Thies, L. Peart-Vissers, P. Boisseau, M. Vincent, A. M. Grabrucker, C. Dubourg, N. Undiagnosed Diseases, W. H. Tan, N. E. Verbeek, M. Granzow, G. W. E. Santen, J. Shendure, B. Isidor, L. Pasquier, R. Redon, Y. Yang, M. W. State, T. Kleefstra, B. Cogne, H. Gem, S. Deciphering Developmental Disorders, S. Petrovski, K. Retterer, E. E. Eichler, J. A. Rosenfeld, P. B. Agrawal, S. Bezieau, S. Odent, Y. Elgersma and S. Mercier, De Novo Mutations in Protein Kinase Genes CAMK2A and CAMK2B Cause Intellectual Disability, Am J Hum Genet, 2017, 101, 768–788.

8. J. R. Stephenson, X. Wang, T. L. Perfitt, W. P. Parrish, B. C. Shonesy, C. R. Marks, D. P. Mortlock, T. Nakagawa, J. S. Sutcliffe and R. J. Colbran, A Novel Human CAMK2A Mutation Disrupts Dendritic Morphology and Synaptic Transmission, and Causes ASD-Related Behaviors, The Journal of Neuroscience, 2017, 37, 2216–2233.

9. M. Proietti Onori, B. Koopal, D. B. Everman, J. D. Worthington, J. R. Jones, M. A. Ploeg, E. Mientjes, B. W. van Bon, T. Kleefstra, H. Schulman, S. A. Kushner, S. Küry, Y. Elgersma and G. M. van Woerden, The intellectual disability-associated CAMK2G p.Arg292Pro mutation acts as a pathogenic gain-of-function, Hum Mutat, 2018, 39, 2008–2024.

10. P. M. F. Rigter, C. de Konink, M. J. Dunn, M. Proietti Onori, J. B. Humberson, M. Thomas, C. Barnes, C. E. Prada, K. N. Weaver, T. D. Ryan, O. Caluseriu, J. Conway, E. Calamaro, C. T. Fong, W. Wuyts, M. Meuwissen, E. Hordijk, C. N. Jonkers, L. Anderson, B. Yuseinova, S. Polonia, D. Beysen, Z. Stark, E. Savva, C. Poulton, F. McKenzie, E. Bhoj, C. P. Bupp, S. Bézieau, S. Mercier, A. Blevins, I. M. Wentzensen, F. Xia, J. A. Rosenfeld, T. C. Hsieh, P. M. Krawitz, M. Elbracht, D. C. M. Veenma, H. Schulman, M. M. Stratton, S. Küry and G. M. van Woerden, Role of CAMK2D in neurodevelopment and associated conditions, Am J Hum Genet, 2024, 111, 364–382.

11. S. J. Coultrap, R. S. Vest, N. M. Ashpole, A. Hudmon and K. U. Bayer, CaMKII in cerebral ischemia, Acta Pharmacol Sin, 2011, 32, 861–872.

12. R. S. Vest, H. O’Leary, S. J. Coultrap, M. S. Kindy and K. U. Bayer, Effective post-insult neuroprotection by a novel Ca(2+)/ calmodulin-dependent protein kinase II (CaMKII) inhibitor, J Biol Chem, 2010, 285, 20675–20682.

13. O. S. Rosenberg, S. Deindl, L. R. Comolli, A. Hoelz, K. H. Downing, A. C. Nairn and J. Kuriyan, Oligomerization states of the association domain and the holoenyzme of Ca^2+^/CaM kinase II, FEBS J., 2006, 273, 682–694.

14. R. Sloutsky, N. Dziedzic, M. J. Dunn, R. M. Bates, A. P. Torres-Ocampo, S. Boopathy, B. Page, J. G. Weeks, L. H. Chao and M. M. Stratton, Heterogeneity in human hippocampal CaMKII transcripts reveals allosteric hub-dependent regulation, Sci Signal, 2020, 13.

15. I. Lučić, L. Héluin, P.-L. Jiang, A. G. Castro Scalise, C. Wang, A. Franz, F. Heyd, M. C. Wahl, F. Liu and A. J. R. Plested, CaMKII autophosphorylation can occur between holoenzymes without subunit exchange, eLife, 2023, 12, e86090.

16. J. Lee, X. Chen and R. A. Nicoll, Synaptic memory survives molecular turnover, Proc Natl Acad Sci U S A, 2022, 119, e2211572119.

17. M. Stratton, I. H. Lee, M. Bhattacharyya, S. M. Christensen, L. H. Chao, H. Schulman, J. T. Groves and J. Kuriyan, Activation-triggered subunit exchange between CaMKII holoenzymes facilitates the spread of kinase activity, Elife, 2014, 3, e01610.

18. E. D. McSpadden, Z. Xia, C. C. Chi, A. C. Susa, N. H. Shah, C. L. Gee, E. R. Williams and J. Kuriyan, Variation in assembly stoichiometry in non-metazoan homologs of the hub domain of Ca(2+) /calmodulin-dependent protein kinase II, Protein Sci, 2019, 28, 1071–1082.

19. Y. Tian, M. A. Shehata, S. J. Gauger, C. Veronesi, L. Hamborg, L. Thiesen, J. Bruus-Jensen, J. S. Royssen, U. Leurs, A. S. G. Larsen, J. Krall, S. M. Ø. Solbak, P. Wellendorph and B. Frolund, Exploring the NCS-382 Scaffold for CaMKIIα Modulation: Synthesis, Biochemical Pharmacology, and Biophysical Characterization of Ph-HTBA as a Novel High-Affinity Brain-Penetrant Stabilizer of the CaMKIIα Hub Domain, J Med Chem, 2022, 65, 15066–15084.

20. P. Wellendorph, S. Høg, C. Skonberg and H. Bräuner-Osborne, Phenylacetic acids and the structurally related non-steroidal anti-inflammatory drug diclofenac bind to specific γ-hydroxybutyric acid sites in rat brain, Fundamental & Clinical Pharmacology, 2009, 23, 207–213.

21. J. Krall, F. Bavo, C. B. Falk-Petersen, C. H. Jensen, J. O. Nielsen, Y. Tian, V. Anglani, K. T. Kongstad, L. Piilgaard, B. Nielsen, D. E. Gloriam, J. Kehler, A. A. Jensen, K. Harpsoe, P. Wellendorph and B. Frolund, Discovery of 2-(Imidazo[1,2-b]pyridazin-2-yl)acetic Acid as a New Class of Ligands Selective for the gamma-Hydroxybutyric Acid (GHB) High-Affinity Binding Sites, J Med Chem, 2019, 62, 2798–2813.

22. Y. Tian, M. A. Shehata, S. J. Gauger, C. K. L. Ng, S. Solbak, L. Thiesen, J. Bruus-Jensen, J. Krall, C. Bundgaard, K. M. Gibson, P. Wellendorph and B. Frolund, Discovery and Optimization of 5-Hydroxy-Diclofenac toward a New Class of Ligands with Nanomolar Affinity for the CaMKIIα Hub Domain, J Med Chem, 2022, 65, 6656–6676.

23. A. P. Torres-Ocampo, C. Ozden, A. Hommer, A. Gardella, E. Lapinskas, A. Samkutty, E. Esposito, S. C. Garman and M. M. Stratton, Characterization of CaMKIIα holoenzyme stability, Protein Sci, 2020, 29, 1524–1534.

24. N. Otmakhov and J. Lisman, Measuring CaMKII concentration in dendritic spines, J Neurosci Methods, 2012, 203, 106–114.

25. M. Bhattacharyya, M. M. Stratton, C. C. Going, E. D. McSpadden, Y. Huang, A. C. Susa, A. Elleman, Y. M. Cao, N. Pappireddi, P. Burkhardt, C. L. Gee, T. Barros, H. Schulman, E. R. Williams and J. Kuriyan, Molecular mechanism of activation-triggered subunit exchange in Ca(2+)/calmodulin-dependent protein kinase II, Elife, 2016, 5.

26. J. B. Myers, V. Zaegel, S. J. Coultrap, A. P. Miller, K. U. Bayer and S. L. Reichow, The CaMKII holoenzyme structure in activation-competent conformations, Nat Commun, 2017, 8, 15742.

27. D. Karandur, M. Bhattacharyya, Z. Xia, Y. K. Lee, S. Muratcioglu, D. McAffee, E. D. McSpadden, B. Qiu, J. T. Groves, E. R. Williams and J. Kuriyan, Breakage of the oligomeric CaMKII hub by the regulatory segment of the kinase, Elife, 2020, 9.

28. M. M. Stratton, Start spreading the news! CaMKII shares activity with naive molecules, Proc Natl Acad Sci U S A, 2022, 119, e2216529119.

29. A. Hudmon, J. Aronowski, S. J. Kolb and M. N. Waxham, Inactivation and self-association of Ca2+/calmodulin-dependent protein kinase II during autophosphorylation, J Biol Chem, 1996, 271, 8800–8808.

30. A. Hudmon, H. Schulman, J. Kim, J. M. Maltez, R. W. Tsien and G. S. Pitt, CaMKII tethers to L-type Ca2+ channels, establishing a local and dedicated integrator of Ca2+ signals for facilitation, J Cell Biol, 2005, 171, 537–547.

31. K. Barcomb, D. J. Goodell, D. B. Arnold and K. U. Bayer, Live imaging of endogenous Ca(2)(+)/calmodulin-dependent protein kinase II in neurons reveals that ischemia-related aggregation does not require kinase activity, J Neurochem, 2015, 135, 666–673.

32. A. Hoelz, A. C. Nairn and J. Kuriyan, Crystal structure of a tetradecameric assembly of the association domain of Ca2+/calmodulin-dependent kinase II, Mol Cell, 2003, 11, 1241–1251.

33. W. Kabsch, Xds, Acta Crystallogr D Biol Crystallogr, 2010, 66, 125–132.

34. P. R. Evans and G. N. Murshudov, How good are my data and what is the resolution?, Acta Crystallogr D Biol Crystallogr, 2013, 69, 1204–1214.

35. J. Agirre, M. Atanasova, H. Bagdonas, C. B. Ballard, A. Basle, J. Beilsten-Edmands, R. J. Borges, D. G. Brown, J. J. Burgos-Marmol, J. M. Berrisford, P. S. Bond, I. Caballero, L. Catapano, G. Chojnowski, A. G. Cook, K. D. Cowtan, T. I. Croll, J. E. Debreczeni, N. E. Devenish, E. J. Dodson, T. R. Drevon, P. Emsley, G. Evans, P. R. Evans, M. Fando, J. Foadi, L. Fuentes-Montero, E. F. Garman, M. Gerstel, R. J. Gildea, K. Hatti, M. L. Hekkelman, P. Heuser, S. W. Hoh, M. A. Hough, H. T. Jenkins, E. Jimenez, R. P. Joosten, R. M. Keegan, N. Keep, E. B. Krissinel, P. Kolenko, O. Kovalevskiy, V. S. Lamzin, D. M. Lawson, A. A. Lebedev, A. G. W. Leslie, B. Lohkamp, F. Long, M. Maly, A. J. McCoy, S. J. McNicholas, A. Medina, C. Millan, J. W. Murray, G. N. Murshudov, R. A. Nicholls, M. E. M. Noble, R. Oeffner, N. S. Pannu, J. M. Parkhurst, N. Pearce, J. Pereira, A. Perrakis, H. R. Powell, R. J. Read, D. J. Rigden, W. Rochira, M. Sammito, F. Sanchez Rodriguez, G. M. Sheldrick, K. L. Shelley, F. Simkovic, A. J. Simpkin, P. Skubak, E. Sobolev, R. A. Steiner, K. Stevenson, I. Tews, J. M. H. Thomas, A. Thorn, J. T. Valls, V. Uski, I. Uson, A. Vagin, S. Velankar, M. Vollmar, H. Walden, D. Waterman, K. S. Wilson, M. D. Winn, G. Winter, M. Wojdyr and K. Yamashita, The CCP4 suite: integrative software for macromolecular crystallography, Acta Crystallographica Section D, 2023, 79, 449–461.

36. A. J. McCoy, R. W. Grosse-Kunstleve, P. D. Adams, M. D. Winn, L. C. Storoni and R. J. Read, Phaser crystallographic software, Journal of Applied Crystallography, 2007, 40, 658–674.

37. D. Liebschner, P. V. Afonine, M. L. Baker, G. Bunkoczi, V. B. Chen, T. I. Croll, B. Hintze, L. W. Hung, S. Jain, A. J. McCoy, N. W. Moriarty, R. D. Oeffner, B. K. Poon, M. G. Prisant, R. J. Read, J. S. Richardson, D. C. Richardson, M. D. Sammito, O. V. Sobolev, D. H. Stockwell, T. C. Terwilliger, A. G. Urzhumtsev, L. L. Videau, C. J. Williams and P. D. Adams, Macromolecular structure determination using X-rays, neutrons and electrons: recent developments in Phenix, Acta Crystallogr D Struct Biol, 2019, 75, 861–877.

38. P. Emsley, B. Lohkamp, W. G. Scott and K. Cowtan, Features and development of Coot, Acta Crystallogr D Biol Crystallogr, 2010, 66, 486–501.

39. N. W. Moriarty, R. W. Grosse-Kunstleve and P. D. Adams, electronic Ligand Builder and Optimization Workbench (eLBOW): a tool for ligand coordinate and restraint generation, Acta Crystallogr D Biol Crystallogr, 2009, 65, 1074–1080.

40. C. E. Blanchet, A. Spilotros, F. Schwemmer, M. A. Graewert, A. Kikhney, C. M. Jeffries, D. Franke, D. Mark, R. Zengerle, F. Cipriani, S. Fiedler, M. Roessle and D. I. Svergun, Versatile sample environments and automation for biological solution X-ray scattering experiments at the P12 beamline (PETRA III, DESY), J Appl Crystallogr, 2015, 48, 431–443.

41. J. B. Hopkins, R. E. Gillilan and S. Skou, BioXTAS RAW: improvements to a free open-source program for small-angle X-ray scattering data reduction and analysis, Journal of Applied Crystallography, 2017, 50, 1545–1553.

42. D. Franke, A. G. Kikhney and D. I. Svergun, Automated acquisition and analysis of small angle X-ray scattering data, *Nuclear Instruments and Methods in Physics Research Section A: Accelerators, Spectrometers*, Detectors and Associated Equipment, 2012, 689, 52–59.

43. K. Manalastas-Cantos, P. V. Konarev, N. R. Hajizadeh, A. G. Kikhney, M. V. Petoukhov, D. S. Molodenskiy, A. Panjkovich, H. D. T. Mertens, A. Gruzinov, C. Borges, C. M. Jeffries, D. I. Svergun and D. Franke, ATSAS 3.0: expanded functionality and new tools for small-angle scattering data analysis, Journal of Applied Crystallography, 2021, 54, 343–355.

44. D. Franke, M. V. Petoukhov, P. V. Konarev, A. Panjkovich, A. Tuukkanen, H. D. T. Mertens, A. G. Kikhney, N. R. Hajizadeh, J. M. Franklin, C. M. Jeffries and D. I. Svergun, ATSAS 2.8: a comprehensive data analysis suite for small-angle scattering from macromolecular solutions, Journal of Applied Crystallography, 2017, 50, 1212–1225.

45. P. V. Konarev, V. V. Volkov, A. V. Sokolova, M. H. J. Koch and D. I. Svergun, PRIMUS: a Windows PC-based system for small-angle scattering data analysis, Journal of Applied Crystallography, 2003, 36, 1277–1282.

46. 46. L. Schrodinger, The PyMOL Molecular Graphics System, 2010, Version 2.0

47. A. G. Kikhney, C. R. Borges, D. S. Molodenskiy, C. M. Jeffries and D. I. Svergun, SASBDB: Towards an automatically curated and validated repository for biological scattering data, Protein Science, 2020, 29, 66–75.

