## Supplementary data for "Ligand-induced CaMKIIα hub Trp403 flip, hub domain stacking and kinase inhibition"

^a^Department of Drug Design and Pharmacology, Faculty of Health and Medical Sciences, University of Copenhagen, Universitetsparken 2, DK-2100 Copenhagen, Denmark; ^b^Department of Biochemistry and Molecular Biology, University of Massachusetts Amherst, MA 01003; ^c^Chemistry-Biology Interface Training Program, University of Massachusetts, Amherst, MA 01003, USA; ^d^HHMI, University of California, Berkeley, CA 94720; ^e^Department of Molecular and Cell Biology, University of California, Berkeley, CA 94720; ^f^California Institute for Quantitative Biosciences, University of California, Berkeley, CA 94720; *^g^*Department of Biochemistry, Vanderbilt University School of Medicine, Nashville, TN 37232; *^h^*Department of Chemistry, University of California, Berkeley, CA 94720; *^i^*Physical Biosciences Division, Lawrence Berkeley National Laboratory, Berkeley, CA 94720.

^*^Main corresponding author: Sara M. Ø. Solbak,; corresponding author on biochemistry and pharmacology: Petrine Wellendorph,; corresponding author on SAXS: Annette E. Langkilde,; corresponding author on X-ray crystallography: Jette S. Kastrup,.

**Table of Contents**

**Table S1**: Data collection and refinement statistics for the crystal structure of CaMKIIα 6x hub with PIPA.

**Table S2**: Volume fractions from the OLIGOMER fits shown in Fig. S5B.

**Table S3.** SAXS data collection details on the CaMKIIα hub domains without and with compounds.

**Figure S1**: Binding of PIPA to CaMKIIα WT and CaMKIIα 6x hub.

**Figure S2**. Thermal shift assay on CaMKIIα hub domains.

**Figure S3**. Inhibition of CaMKIIα syntide-2 phosphorylation.

**Figure S4**. SAXS on CaMKIIα hub domains.

**Figure S5**. Models of self-association and corresponding fits to SAXS data.

**Figure S6**. MP histograms of CaMKIIɑ WT hub and CaMKIIɑ W403L hub.

**Figure S7**. Comparison of CaMKIIα 6x hub structures with PIPA and 5-HDC.

**Figure S8**. Crystal packing of tetradecameric CaMKIIα 6x hub.

**Table S1.** Data collection and refinement statistics for the crystal structure of CaMKIIα 6x hub with PIPA.

| **Crystal data** | |
| --- | --- |
| Space group | *C 2 2 2_1_* |
| Unit cell axes *a, b*, *c* (Å) | 103.05, 182.92, 107.76 |
| Unit cell axes *α, β*, γ (°) | 90, 90, 90 |
| Molecules in a.u.^a^ | 7 |
| **Data collection** | |
| Beamline | ALS beamline 8.2.2 |
| Wavelength (Å) | 1.0000 |
| Resolution (Å) | 46.48-2.10 (2.16-2.10)^b^ |
| No. of unique reflections | 59,633 (4,582)^b^ |
| Average multiplicity | 14.8 (15.0)^b^ |
| Completeness (%) | 100 (100)^b^ |
| *R*_pim_ (%) | 2.9 (63.7)^b^ |
| Mean I/σ(I) | 14.4 (1.3)^b^ |
| Wilson B (Å^2^) | 48.3 |
| CC_1/2_ | 1 (0.58)^b^ |
| **Refinement** | |
| Amino-acid residues built (chain A/B/C/D/E/F/G) | 132/133/132/132/133/132/130 |
| PIPA/acetate/PEG/water | 2/6/1/102 |
| *R*_work_/*R*_free_ (%) | 22.4/25.1^c^ |
| *Average B values (Å^2^) for:* | |
| Amino-acid residues (chain A/B/C/D/E/F/G) | 85.1/86.9/94.8/99.8/77.2/70.4/72.9 |
| PIPA/acetate/PEG/water | 77.2/60.0/91.9/57.7 |
| RMSD bond length (Å)/angles (°) | 0.003/0.64 |
| Ramachandran outliers/favored (%) | 0.0/97.6^d^ |
| Rotamer outliers (%) /Cβ outliers (%) /Clash score | 0.3/0/6.6^d^ |

^a^ a.u: asymmetric unit of the crystal

^b^ Outershell values are shown in parentheses

^c^ R_free_ is equivalent to R_work_, but calculated with 5 % of reflections omitted from the refinement process

^d^ MolProbity statistics from phenix

**Table S2**: Volume fractions from the OLIGOMER fits shown in Fig. S5B. Volume fractions larger than 2 SD are highlighted in bold. The models are listed by number of dodecamers, e.g., d4 consists of four stacked dodecamers.

|  | WT hub  + PIPA (i) | WT hub  + PIPA (ii) | WT hub  + PIPA (iii) | WT hub  + PIPA (iv) | WT hub  + PIPA (ii)  remeasure |
| --- | --- | --- | --- | --- | --- |
| **χ^2^** | 1.29 | 5.62 | 5.94 | 2.45 | 4.43 |
| **dodecamer** | **0.820±0.007** | **0.682±0.003** | **0.623±0.003** | **0.367±0.002** | **0.717±0.002** |
| **d2** | **0.097±0.014** | **0.184±0.006** | **0.095±0.005** | **0.149±0.007** | 0 |
| **d3** | 0 | 0 | 0.031±0.007 | 0.102±0.007 | 0 |
| **d4** | 0 | 0 | 0 | 0. | 0 |
| **d5** | 0 | 0 | 0 | 0 | 0 |
| **d6** | 0 | 0 | 0 | 0 | 0 |
| **d7** | 0 | 0 | 0 | **0.053±0.020** | 0 |
| **d8** | 0 | 0 | 0 | 0 | 0 |
| **d9** | 0 | 0 | 0 | 0.017±0.032 | 0 |
| **d10** | 0 | 0 | 0 | 0 | 0 |
| **d11** | 0 | 0 | 0 | **0.234±0.031** | 0 |
| **d12** | 0 | 0 | 0 | 0 | 0 |
| **d13** | 0 | 0.043±0.023 | **0.229±0.018** | 0.033±0.019 | 0 |
| **d14** | 0 | 0 | 0 | 0 | 0 |
| **d15** | 0 | **0.078±0.023** | 0.023±0.017 | 0 | 0 |
| **d16** | **0.064±0.023** | 0 | 0 | 0 | 0 |
| **d20** | 0.040±0.023 | 0.012±0.007 | 0 | 0 | 0 |
| **d24** | 0.005±0.015 | 0 | 0 | 0 | 0 |
| **d32** | 0 | 0 | 0 | 0.001±0.005 | 0 |
| **d40** | 0 | 0 | 0 | **0.044±0.005** | **0.283±0.001** |

**Table S3.** SAXS data collection details on the CaMKIIα hub domains without and with compounds.

| Sample  (all hub domains) | Conc. (mg/mL) | Facility | Wavelength (Å) | Detector distance (m) | Exposure time | Measured q range (Å^-1^)* |
| --- | --- | --- | --- | --- | --- | --- |
| WT | 6.1 | P12 | 1.2398 | 3.0 | 40 x 0.090 s | 0.00224-0.734 |
| WT | 4.6 | CPHSAXS | 1.3428 | 1.0355 (S) | 10 x 60 s (S) | 0.01173-0.306 |
| WT | 2.3 | CPHSAXS | 1.3428 | 1.0355 (S) | 10 x 60 s (S) | 0.01173-0.306 |
| WT | 1.1 | CPHSAXS | 1.3428 | 1.0355 (S) | 10 x 60 s (S) | 0.01173-0.306 |
| WT  + HOCPCA | 2.9 | CPHSAXS | 1.3428 | 0.610 (M)  1.0355 (S)  1.4355 (E) | 10 x 60 s (M) 10 x 60 s (S)  10 x 120 s (E) | 0.00559-0.306 |
| WT  + 5-HDC | 3.4 | P12 | 1.2398 | 3.0 | 40 x 0.09 s | 0.00224-0.734 |
| WT  + PIPA (i) | 1.2 | CPHSAXS | 1.3402 | 0.6325 (M)  1.4558 (E) | 40 x 60 s (M)  60 x 120 s (E) | 0.0027-0.508 |
| WT  + PIPA (ii)** | 2.4 | CPHSAXS | 1.3404 | 0.6325 (M) | 40 x 60 s (M) | 0.0159-0.513 |
| WT  + PIPA (ii)**  Remeasure after 12 h | 2.4 | CPHSAXS | 1.3404 | 0.6325 (M) | 40 x 60 s (M) | 0.0159-0.513 |
| WT  + PIPA (iii)** | 3.9 | CPHSAXS | 1.3404 | 0.6325 (M) | 40 x 60 s (M) | 0.0159-0.513 |
| WT  + PIPA (iv)*** | 2.3 | P12 | 1.2400 | 3.0 | 30 x 0.095 s | 0.00205-0.447 |
| 6x | 3.4 | P12 | 1.2398 | 3.0 | 40 x 0.090 s | 0.00224-0.734 |
| 6x | 4.2 | CPHSAXS | 1.3428 | 1.0355 (S) | 10 x 60 s (S) | 0.01173-0.306 |
| 6x | 2.1 | CPHSAXS | 1.3428 | 1.0355 (S) | 10 x 60 s (S) | 0.01173-0.306 |
| 6x | 1.1 | CPHSAXS | 1.3428 | 1.0355 (S) | 10 x 60 s (S) | 0.01173-0.306 |
| 6x  + 5-HDC | 4.4 | CPHSAXS | 1.3428 | 1.0355 (S)  1.4355 (E) | 10 x 60 s (S)  10 x 120 s (E) | 0.00559-0.306 |
| 6x  + PIPA | 6.7 | CPHSAXS | 1.3402 | 0.6325 (M)  1.4558 (E) | 40 x 60 s (M)  60 x 120 s (E) | 0.0027-0.508 |
| W403L | 6.4 | P12 | 1.2400 | 3.0 | 30 x 0.095 s | 0.00205-0.447 |
| W403L  + 5-HDC | 6.0 | CPHSAXS | 1.3402 | 0.6325 (M)  1.4558 (E) | 40 x 60 s (M)  60 x 120 s (E) | 0.0027-0.508 |
| W403L  + PIPA | 6.8 | P12 | 1.2400 | 3.0 | 30 x 0.095 s | 0.00205-0.447 |

*Data collected. Subsequently cropped with q_min_ according to Guinier analyses and q_max_ for additional analyses set to ≤0.4 Å^-1^.

**Some sample development / heterogeneity during data collection, thus only a full average from the M setting used.

***Soluble fraction only, i.e., supernatant after sample with visible aggregation was centrifuged.

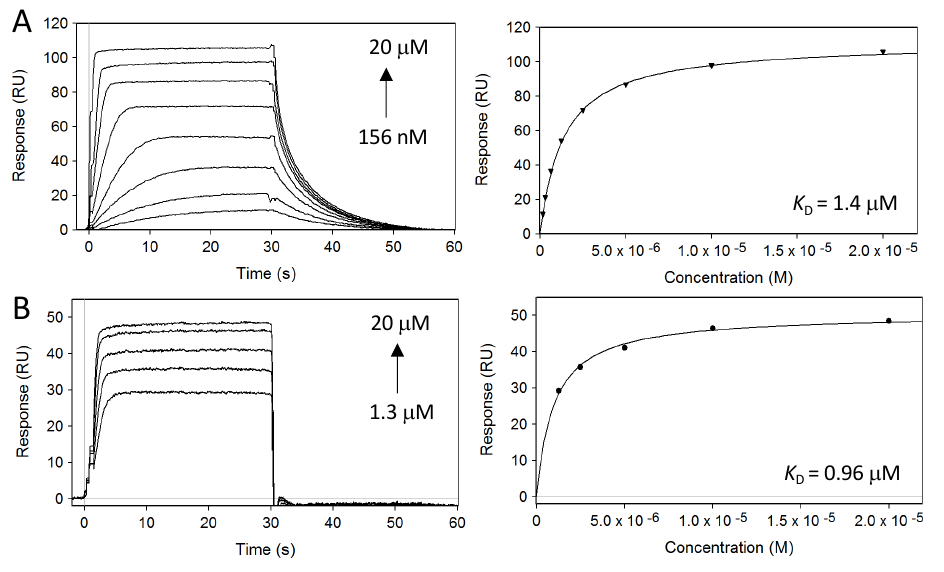

**Figure S1. Binding of PIPA to CaMKIIα 6x hub and CaMKIIα holenzyme. A)** Concentration-dependent binding of PIPA to CaMKIIα 6x hub measured by SPR (left) and SPR Langmuir-binding isotherm (right). **B)** Concentration-dependent binding of PIPA to immobilized CaMKIIα holoenzyme measured by surface plasmon resonance (left) and SPR Langmuir-binding isotherm (right).

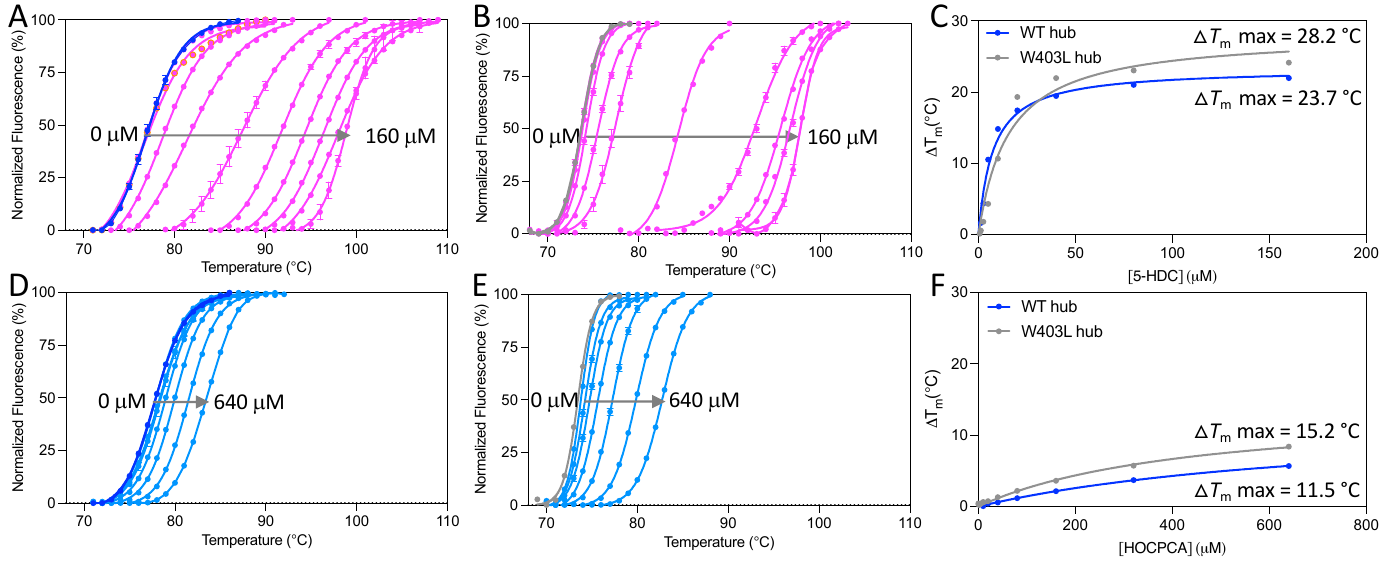

**Figure S2. Thermal shift assay on CaMKIIα hub domains. A)** Right-shifted thermal shift assay melting curves of CaMKIIα WT hub upon binding of HOCPCA. **B)** Right-shifted thermal shift assay melting curves of CaMKIIα W403L hub upon binding of HOCPCA. **C)** Thermal melting point of CaMKIIα WT hub (blue) and CaMKIIα W403L hub (grey) plotted against increasing concentrations of HOCPCA (representative data of n = 3). **D)** Right-shifted thermal shift assay melting curves of CaMKIIα WT hub upon binding of 5-HDC. **E)** Right-shifted thermal shift assay melting curves of CaMKIIα W403L hub upon binding of 5-HDC. **F)** Thermal melting point of CaMKIIα WT hub (blue) and CaMKIIα W403L hub (grey) plotted against increasing concentrations of 5-HDC (representative data of n = 3).

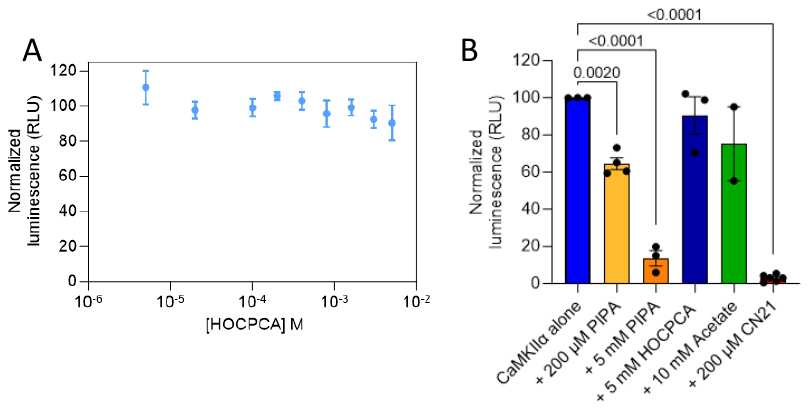

**Figure S3.** **Inhibition of CaMKIIα syntide-2 phosphorylation.** **A)** No concentration-dependent inhibition of CaMKIIα syntide-2 phosphorylation by HOCPCA, investigated with the luminescence-based ADP-Glo kinase assay (n = 3, mean ± SEM). **B)** Comparison of kinase inhibition by PIPA (200 µM and 5 mM), HOCPCA (5 mM), acetate (10 mM), and CN21 (200 µM) (n = 2-4, mean ± SEM).

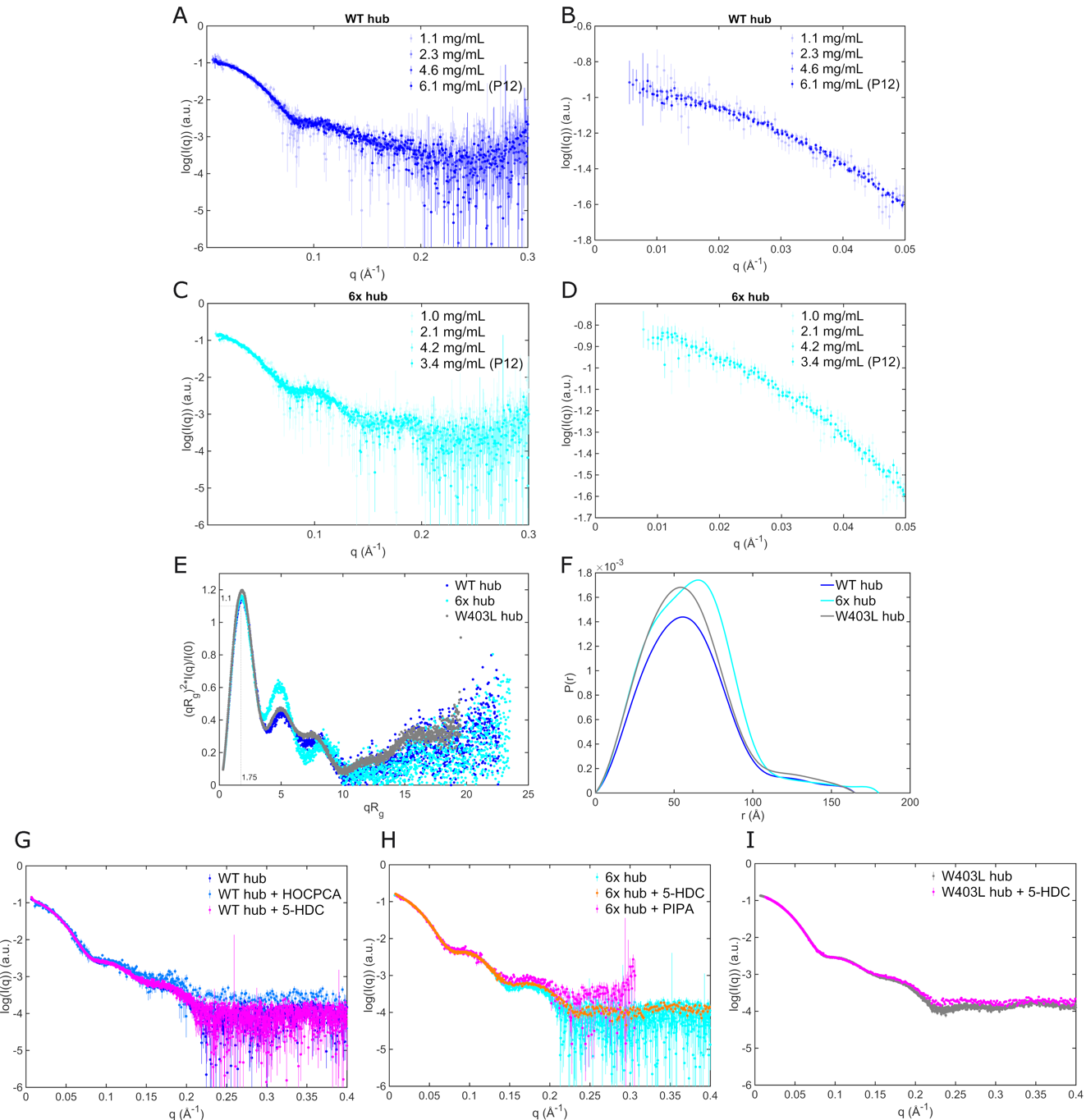

**Figure S4. SAXS on CaMKIIα hub domains. A-B)** Scattering curves for CaMKIIα WT hub at different concentrations and from both lab source (unmarked) and synchrotron (P12). **C-D)** Scattering curves of CaMKIIα 6x hub at different concentrations and from both lab source (unmarked) and synchrotron (P12). **E)** Kratky plots of CaMKIIα WT, 6x, and W403L hub domains. **F)** CaMKIIα WT, 6x, and W403L hub domains pair distance distribution functions. **G)** CaMKIIα WT hub without and with compounds (1000 mM HOCPCA and 100 mM 5-HDC). **H)** Scattering curves of CaMKIIα 6x hub without and with compounds (200 mM PIPA and 100 mM 5-HDC). **I)** Scattering curves of CaMKIIα W403L hub without and with 100 mM 5HDC.

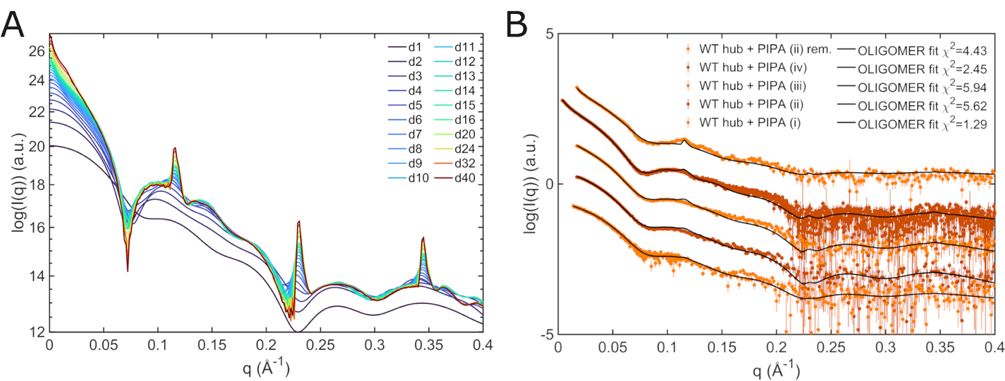

**Figure S5. Models of self-association and corresponding fits to SAXS data. A**) Theoretical scattering curves for stacked dodecamers of CaMKIIα hub domain structures used in the OLIGOMER pool. The curves are ordered by number of dodecamers in the respective model, e.g., d4 consists of four stacked dodecamers. All models are based on PDB entry 5IG3, as illustrated in Fig. 3E. **B)** Fits to SAXS data of CaMKIIα WT hub with PIPAobtained from OLIGOMER with different distributions of self-associated stacks (cf Table S2). The scattering curves have been translated for clarity.

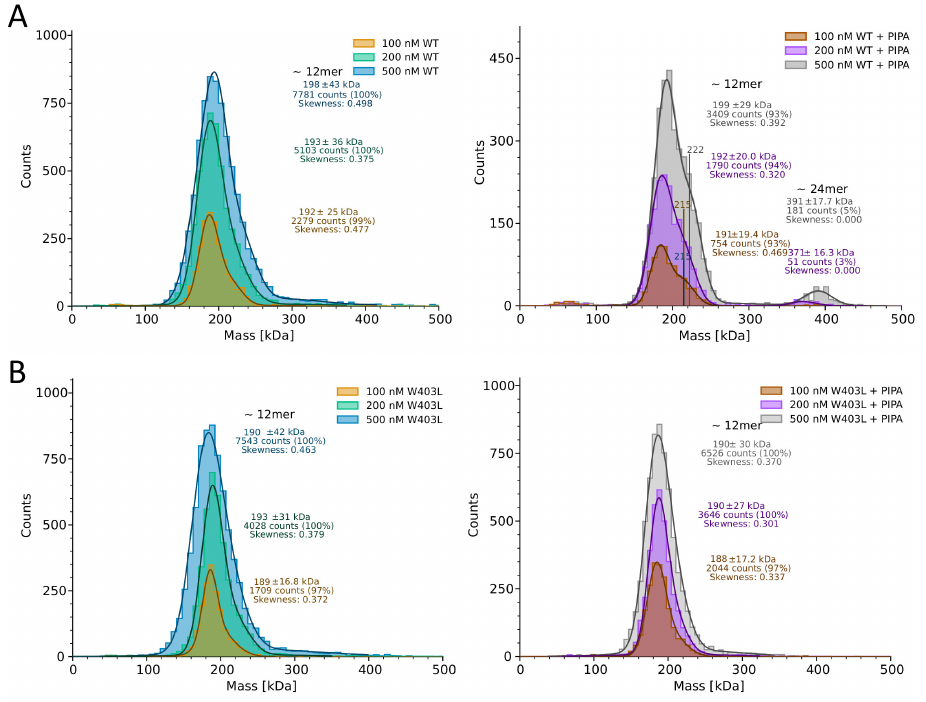

**Figure S6.** **MP histograms of CaMKIIɑ WT hub and CaMKIIɑ W403L hub. A**) Representative MP histograms showing CaMKIIɑ WT hub diluted to 500 nM, 200 nM and 100 nM without (left) and with the addition of PIPA (right). **B)** Representative MP histograms of CaMKIIɑ W403L hub diluted to 500 nM, 200 nM, and 100 nM without (left) and with the addition of PIPA (right). Each count indicates a single molecule.

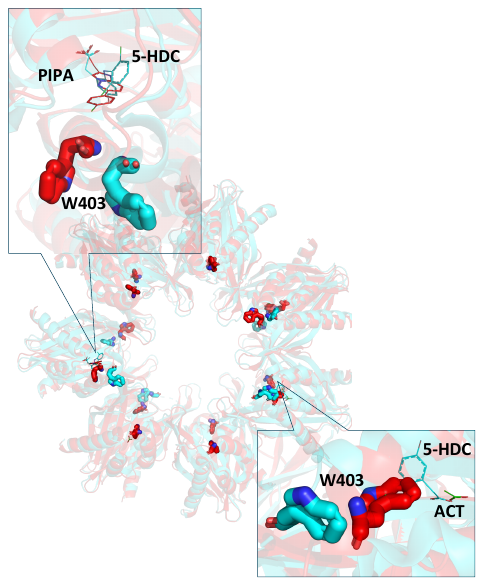

**Figure S7.** **Comparison of CaMKIIα 6x hub structures with PIPA and 5-HDC**. Superimposed X-ray structures (PDB entry 7REC) of tetradecameric CaMKIIα 6x hub (14-mer generated from 7-mer) with PIPA (red) and 5-HDC (cyan). The proteins are shown as cartoon. Zoomed-in (top left) view shows the binding site of CaMKIIα 6x hub compounds (PIPA and 5-HDC) as lines and the corresponding Trp403 positioned outwards in sticks representation. Zoomed-in (bottom right) view shows the binding site of CaMKIIα 6x hub compounds (acetate (red) and 5-HDC) as lines and the corresponding Trp403 flipped inward (acetate)/outward (5-HDC).

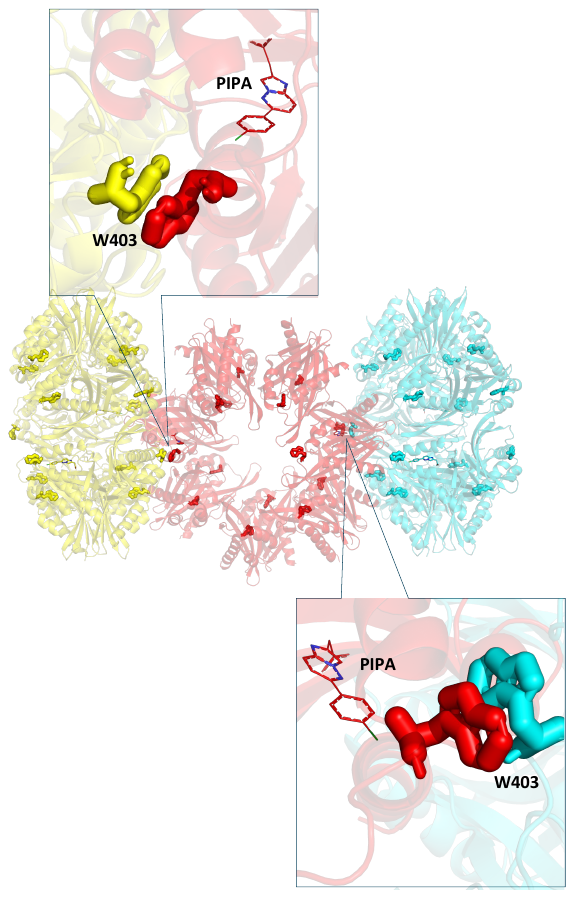

**Figure S8. Crystal packing of tetradecameric CaMKIIα 6x hub (14-mer generated from 7-mer) shown as cartoon**. Zoomed-in (top left and bottom right) views show the binding site for CaMKIIα 6x hub compounds (here PIPA) in red line representation and Trp403 in sticks representation. Trp403 from the corresponding symmetry mates are colored yellow and cyan.
